# Vgll2 and Tead1 Govern Generation of Mouse and Human Hypothalamic Hypocretin (Orexin) Neurons

**DOI:** 10.64898/2026.03.30.715421

**Authors:** Ruofan Wei, Amirhossein Sheikhshahrokh, Elnaz Saeidi, Cecilia Gomez-Inclan, Aswathi Gopalakrishnan, Brad Balderson, Mikael Bodén, Michael Piper, Stefan Thor

## Abstract

The hypothalamic hypocretin (Hcrt; aka orexin) neuropeptide neurons are crucial for wakefulness and their malfunction results in narcolepsy; a devastating disorder without current cures. Transplantation of mouse Hcrt cells into mouse narcolepsy models is pointing to cell therapy as a potential treatment for narcolepsy. However, progress in this area requires decoding the developmental pathways generating human HCRT neurons and a deeper understanding of their diversity and physiology. Here, we identify the hypothalamic progenitor domain generating mouse Hcrt neurons and propose a conserved mouse/human genetic cascade driving Hcrt specification. This cascade involves the Vgll2 and Tead1 transcription factor partners, which we find are crucial for mouse Hcrt neuron generation. Conversely, co-misexpression of VGLL2 and TEAD1 in human stem cell derived hypothalamic organoids can induce HCRT neuron differentiation. These findings provide an important element for the development of human induced HCRT neurons for physiological studies and for cell therapy in narcolepsy.

## INTRODUCTION

The hypothalamus is a key regulator of many bodily functions, including energy and fluid balance, thermoregulation, sleep-wake states, stress responses, growth and reproduction.^1^ The pervasive homeostatic role of the hypothalamus relies upon its staggering neuronal diversity, with extensive neuropeptide expression, intricate neurotransmitter/neuropeptide co-expression, widespread connectome and elaborate neuroendocrine secretion.^2^ Unsurprisingly, single cell/nuclei and spatial transcriptomic analyses has uncovered hundreds of transcriptionally distinct cell types in the mouse, macaque and human hypothalamus, pointing to, arguably, the greatest cell diversity per volume in the mammalian body.^3–38^ While substantial progress has been made regarding the spatio-temporal patterning of the developing hypothalamus,^2,3,31,39–48^ the subsequent genetic pathways dictating the terminal cell identity of the hundreds of identified cell types remain poorly understood.

Viewed from anterior to posterior, the hypothalamus contains four main regions: preoptic, anterior, tuberal and mammillary, with each region further subdivided into several nuclei.^2,46^ The Hypocretin (Hcrt; aka Orexin) neuropeptide neurons are chiefly located in the tuberal region and the lateral hypothalamus nucleus (LH).^49,50^ Hcrt neurons are crucial for maintaining wakefulness, but are also involved in regulating feeding, emotional behaviour and reward seeking.^51–53^ Studies mainly in rodents have revealed that the activity of Hcrt neurons is controlled by a number of other neurotransmitters and circulatory factors,^54^ but the relevance of these findings to human HCRT neurons is less clear. Early single cell RNA sequencing (sc-RNA-seq) analysis pointed to gene expression and putative lineage relatedness between mouse Hcrt neurons and another dorso-tuberal neuropeptide cell group, the Npvf neurons,^17^ involved in gating fertility^55,56^ and circadian motor activity.^57^ Recent sn/sc-RNA-seq studies, involving orders of magnitude more cells, have confirmed the notion of relatedness between Hcrt and Npvf neurons and placed them within a distinct cluster of glutamatergic neurons, denoted C66-18:Rfx4.GLU-4, identified in adult mouse,^5^ or excitatory neuron cluster 21 (EN21), identified during developmental stages in mouse, macaque and human.^30^ We will chiefly refer to this cluster by its developmental definition, EN21.

Hcrt or Hcrt receptor knockout, as well as Hcrt cell loss, by genetic manipulation or autoimmunity, results in narcolepsy in mice, dogs and humans.^58–64^ In humans, narcolepsy is a devastating disease without current cure, affecting ∼1/2,000 individuals.^65^ Transplantation of purified mouse Hcrt-GFP neurons into Hcrt cell-deficient mice resulted in partial rescue of narcoleptic phenotypes,^66,67^ providing proof-of-principle for the development of cell-based therapies for human narcolepsy.^68^ Human induced pluripotent stem cells (hiPSCs) can be driven into hypothalamic-like 2D and 3D (organoid) differentiation.^69^ These cultures can contain HCRT neurons, but as anticipated from their scarcity in the hypothalamus, in low proportions.^70^ The generation of hiPSC-derived HCRT neurons sets the stage for cell transplantation development, pre-clinically and subsequently clinically, but requires a deeper understanding of HCRT cell specification to more readily generate large numbers of these neurons.

Three transcription factors (TFs) have been genetically demonstrated to be important for mouse Hcrt neuron generation: Ebf2, Dbx1 and Lhx9.^8,71–73^ However, none of these three mouse mutants show a complete loss of Hcrt cells. Moreover, gene expression analysis has not identified *Dbx1* or *Ebf2* expression in mature mouse or human Hcrt neurons,^26,29,74^ pointing to that their effect on Hcrt neurons may be a consequent of patterning or lineage defects in the developing hypothalamus. By contrast, *Lhx9* is expressed by mature Hcrt neurons,^26,29^ and ectopic *Lhx9* expression in the developing mouse hypothalamus can trigger extra Hcrt cells.^72^ However, because terminal cell fates is often controlled by combinatorial codes of TFs,^75^ additional TFs are likely involved in the specification of Hcrt neurons. The Vestigial like TF co-factor gene *Vgll2* is expressed in the developing mid-tuberal mouse hypothalamus, generating the ventro-medial hypothalamus (VMH),^12^ but is also selectively expressed in the dorso-medial (DMH) and LH regions, specifically in adult Hcrt and Npvf neurons.^7,8^ VGLL factors (Vgll1/2/3/4 in mammals) physically and genetically act with TEAD TFs (Tead1/2/3/4 in mammals).^76,77^ Gene expression analysis^19^ pointed to Tead1 as the most likely Vgll2 partner in Hcrt and Npvf neurons, suggesting that the Vgll2-Tead1 pair could play important roles in Hcrt and Npvf cell specification.

To address the role of *Vgll2* and *Tead1* in Hcrt and Npvf cell specification, we analysed their expression profiles in the developing mouse hypothalamus, using a combination of newly generated and publicly available expression data. Integrating Vgll2 and Tead1 expression with a number of hypothalamic patterning TFs resulted in the identification of a specific progenitor domain in the developing dorso-tuberal hypothalamus, proposed as pT1-1 (progenitor Tuberal 1-1), and a putative genetic pathway towards the EN21/Hcrt/Npvf neurons. This pathway incorporated the known Hcrt determinants *Dbx1* and *Lhx9* and proposed *Vgll2* and *Tead1* as additional determinants. To functionally test this notion, we generated CNS-conditional mouse mutants for *Vgll2* and *Tead1* (*Vgll2^cKO^*and *Tead1^cKO^*). We observed a complete/near-complete loss of both Hcrt and Npvf neurons in both mutants, evident by immunostaining and sn-RNA-seq analysis. *Vgll2^cKO/+^;Tead1^cKO/+^* trans-heterozygotes show significantly reduced Hcrt and Npvf cell numbers, when compared to either heterozygote, indicating combinatorial involvement. Neither mutant displayed altered gene expression of *Ebf2*, *Dbx1* and *Lhx9*, nor loss of the EN21 neuron cluster, indicating that *Vgll2* and *Tead1* act in parallel to and/or downstream of these regulators during Hcrt/Npvf cell specification. We analysed gene expression for the proposed pT1-1/EN21/Hcrt/Npvf pathway in published human sn/sc-RNA-seq datasets, which suggested a similar pathway to that observed in mouse. To begin testing this notion, we addressed the sufficiency of *VGLL2* and *TEAD1* in human cells, by misexpressing them in human induced pluripotent stem cell (hiPSC) derived hypothalamic organoids. We find that *VGLL2*-*TEAD1* co-misexpression could induce HCRT neurons (iHCRT), which displayed transcriptional profiles related to several human HCRT subtypes. These findings place Hcrt neuron specification within a conserved spatio-temporal developmental framework within the dorso-tuberal hypothalamus, paving the way for hiPSC-derived iHCRT neurons for physiological studies and for the development of cell therapy in narcolepsy.

## RESULTS

### Spatio-temporal development of mouse and human Hcrt and Npvf neurons

To map the developmental trajectory of the EN21 cluster and the Hcrt and Npvf neurons, and place Vgll2 and Tead1 expression within this framework, we conducted immunostaining and sn-RNA-seq analysis and interrogated previously published gene expression data sets.

Allen Brain Atlas (ABA) *in situ* hybridization data,^78^ conducted at E11.5, E13.5, E15.5 and E18.5, revealed onset of *Npvf* expression at E13.5 and *Hcrt* at E15.5, in the developing dorso-lateral region of the tuberal mouse hypothalamus (Figure 1B-C). This expression was also evident from recent spatial transcriptomic analysis at E13.5 (Figure 1H). Spatial transcriptomic analysis at E13.5^30^ confirmed expression of *Vgll2* in the mid- and dorso-lateral part of the tuberal hypothalamus (Figure 1H). ABA *in situ* hybridization and the spatial transcriptomic analysis showed expression of *Tead1* and *Tead2* in the developing mouse hypothalamus, marked by *Rax* expression, with limited, if any expression of *Tead3* and *Tead4* (Figure 1A, 1D, 1H, S1A-C). *Tead1* was expressed both along the ventricular zone, in progenitors, and laterally in differentiating cells, whereas *Tead2* appeared confined to the ventricular zone (Figure 1D, 1H, S1A). *Tead1* expression was evident in the region of *Hcrt* and *Npvf* cells (Figure 1D, 1H). *Vgll1* displayed no clear expression, while *Vgll3* and *Vgll4* displayed limited and scattered expression throughout the hypothalamus (Figure S1D-F).

**Figure 1.**
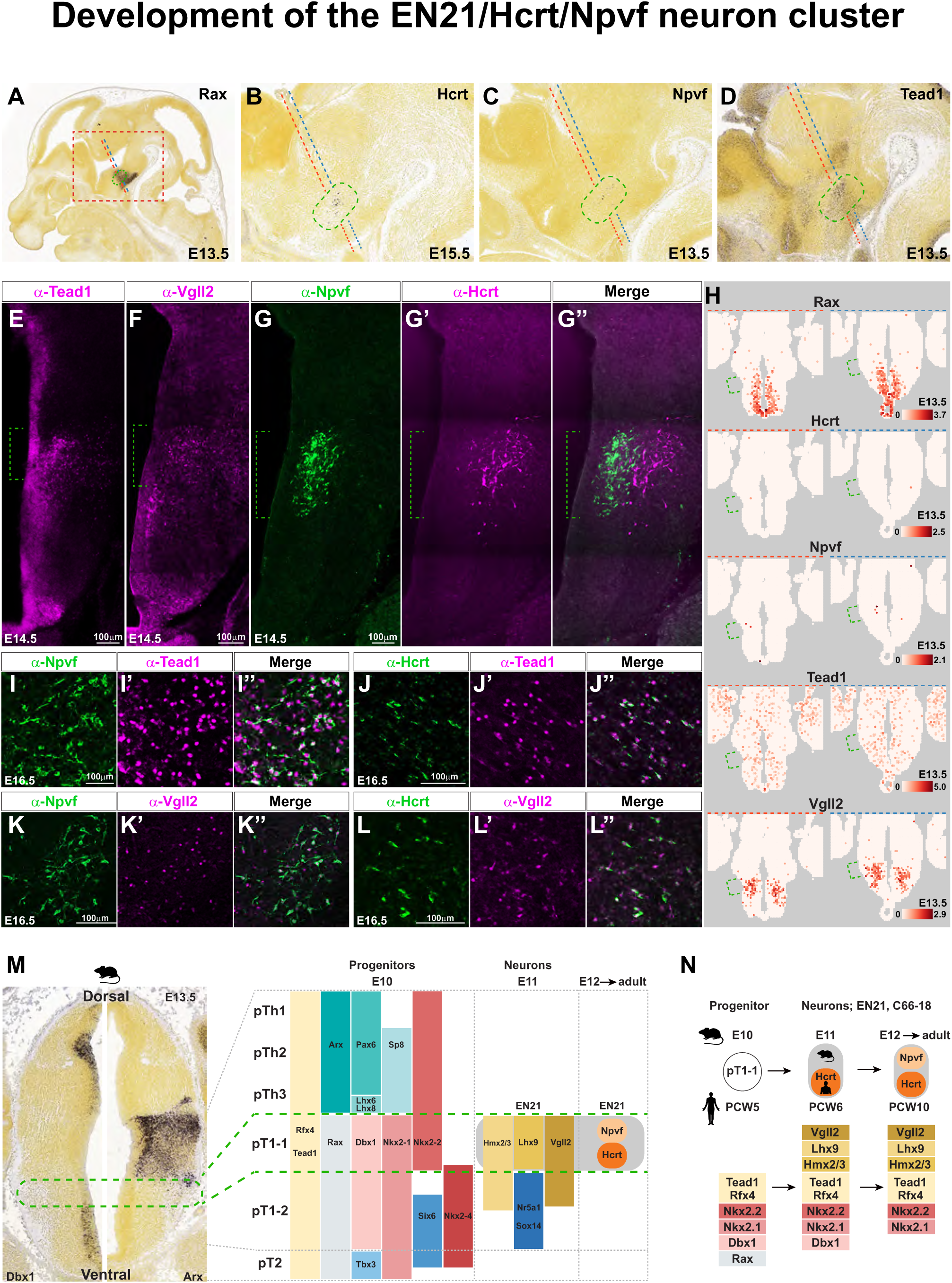
Spatio-temporal development of the mouse hypothalamic pT1-1 progenitor domain and the EN21/Hcrt/Npvf neuronal cluster. **(A-D,H)** Expression of *Rax*, *Hcrt*, *Npvf*, *Tead1* and *Vgll2* and in the developing mouse tuberal hypothalamus based upon sagittal ABA *in situ* hybridization images^78^ or transverse spatial RNA-seq data.^30^ Black dashed box in (A) outlines the regions shown in (B-D). Green brackets and boxes depict the putative pT1-1/EN21 domain. Blue and red dashed lines depict the estimated level of transverse sections in the spatial RNA-seq data when compared to the sagittal *in situ* hybridization images. **(E-G’’)** Expression of Tead1, Vgll2, Hcrt and Npvf in the developing mouse tuberal hypothalamus (brackets depict the putative pT1-1/EN21 domain). **(I-L’’)** Double immunostaining for Hcrt, Npvf, Tead1 and Vgll2, showing that most, if not all Hcrt and Npvf neurons express Tead1 and Vgll2. **(M)** Outline of the development of the tuberal hypothalamus in transverse section, with the proposed pT1-1 progenitor domain (green box) and the emerging EN21 neuronal cluster (see Figures S1-S4 for supporting information). **(N)** Outline of gene expression during development of pT1-1 progenitors and the EN21 cluster, in mouse and human.

Immunostaining confirmed the mRNA expression data, revealing early onset of Tead1 expression along the ventricular zone throughout the hypothalamus and in a subset of differentiating regions, including the dorso-lateral tuberal region (Figure 1E). Vgll2 expression was restricted to the mid- and dorso-lateral tuberal hypothalamus (Figure 1F). Npvf and Hcrt immunostaining agreed with mRNA expression and was detected in the dorso-tuberal region, with robust expression commencing at E14.5 (Figure 1G-G’’). Importantly, Vgll2 and Tead1 are expressed in most, if not all Hcrt and Npvf cells from neuropeptide expression onset and onward to P20 (Figure 1I-L’’).

Previous *in situ* gene expression mapping identified the *Hcrt* domain as located in the dorsal-most *Nkx2-1* region in the tuberal hypothalamus, in a stripe of *Lhx9* expression and dorsally abutted by *Lhx6*, *Lhx8* and *Arx*.^18^ Building upon this mapping we again made use of ABA *in situ* hybridization^78^ and recent spatial transcriptomic data^30^ to further integrate *Tead1*, *Vgll2*, *Hcrt* and *Npvf* expression within the developing hypothalamus. Comparison of gene expression of *Tead1*, *Vgll2*, *Hcrt* and *Npvf* with 17 previously identified TF genes selectively expressed in the hypothalamus and thalamus^2,3,31,39–48^ helped us further refine the dorso-ventral patterning of the tuberal hypothalamus and define the EN21/Hcrt/Npvf domain (Figure 1M, S2A-P). ABA revealed limited, if any expression of *Ebf2* in the hypothalamus at any stage. The spatial transcriptomic data showed some scattered expression in the tuberal hypothalamus, but with little if any overlap with the other EN21/Hcrt/Npvf markers (not shown).

Next, we surveyed gene expression within publicly available mouse embryonic and adult sn/sc-RNA-seq data.^5,30^ This revealed that *Tead1* expression commenced at E10.5 in many *Rax*^+^ progenitors and subsets of neurons (Figure S3A), while *Vgll2* expression commenced at E11.5 in subsets of neurons. In agreement with spatial transcriptomic and *in situ* data, a subset of the aforementioned TF genes was expressed in progenitors from E10.5 and onward (Figure S3C). The EN21 cluster became evident at E11.5 and expressed *Vgll2* and *Tead1*, as well as seven of the other TF genes (Figure S3B, 3D). *Hcrt* and *Npvf* expression became evident by E12.5 (Figure S3D). *Rax* and *Dbx1* expression was not robustly maintained in the EN21 cluster into P0 and adulthood, while the other eight genes; *Vgll2*, *Lhx9*, *Hmx2*, *Hmx3* (*Hmx2/3*), *Tead1*, *Rfx4*, *Nkx2-2* and *Nkx2-1* maintained their expression in EN21, as well as in the adult C66-18:Rfx4.GLU-4 cluster (Figure S3D). *Ebf2* was expressed by a subset of progenitors and intermediate progenitor cells, with some overlap with *Lhx9* in the latter, but not robustly expressed in the EN21/Hcrt/Npvf cells at any stage (not shown). *Vgll2* expression was previously described in the developing VMH i.e., the mid-tuberal region^12^ and the sn/sc-RNA-seq data revealed *Vgll2* expression in the two recently identified DMH glutamatergic cell clusters EN7 and EN8^30^ (Figure S4A-F).

Similar to mouse, the expression of the HCRT and NPVF neuropeptides has been observed in the adult human tuberal hypothalamus, in the LH and DMH for HCRT^60,61,79–81^ and in the DMH for NPVF.^82^ Recently published human developmental sn/sc-RNA-seq data sets^30^ allowed us to mirror the expression analysis conducted in mouse, revealing extensive similarity between mouse and human. Specifically, a subset of TF genes, including *TEAD1*, commenced expression at post-conception week (PCW) 5 in many hypothalamic *RAX*^+^ progenitors (Figure S5A, S5C). *VGLL2*, *LHX9*, *RFX4* and *HMX2/3* expressed in subsets of neurons, including the EN21 cluster, with an onset at PCW6 (Figure S5B, S5D). *HCRT* expression commenced by PCW6, while *NPVF* was not apparent until PCW10 (Figure S5D). Like mouse, *RAX* and *DBX1* expression was only transiently expressed in the EN21 cluster, while the other eight TF genes continued expressing into later stages (Figure S5D).

The comprehensive expression analysis provided by *in situ* hybridization, BacTRAP, sn/sc-RNA-seq, spatial-RNA-seq and antibody staining places the EN21/C66-18 cluster, Vgll2, Tead1, Hcrt and Npvf within an emerging spatio-temporal framework in the tuberal hypothalamus (Figure 1M). First, we propose a sub-division of the previously identified progenitor type pT1^83^ into pT1-1 and pT1-2, defined by the differential expression of several patterning genes, most prominently with pT1-1 being Nkx2-2^+^/Six6^-^ and pT1-2 being Nkx2-2^-^/Six6^+^/Tbx3^-^, and with pT2 defined by being Tbx3^+^. Second, several of the TFs expressed in the pT1-1 progenitor domain continue expressing into the emerging EN21 cell cluster. Third, expression of four genes emerges lateral to the pT1-1 domain, in the differentiation zone: *Hmx2/3*, *Lhx9* and *Vgll2*, helping to define the EN21 cell cluster, which is furthermore defined by being Nr5a1^-^/Sox14^-^. Fourth, expression of Hcrt and Npvf commences. Fifth, while two TF genes, *Rax* and *Dbx1*, are downregulated over time in Hcrt and Npvf cells, the other eight TFs continue expressing into adulthood. The human mapping is more limited in scope, but the previous immunostaining and the sc/sn-RNA-seq data indicates a similar scenario in humans (Figure 1N). Underscoring the relevance of this model is that one patterning gene, *Dbx1*, and one postmitotic gene, *Lhx9*, are both known to be important for mouse Hcrt cell generation.^8,72,73^ Importantly, the expression of *Vgll2* and *Tead1* is also in line with their potential roles in specifying Hcrt and Npvf neurons.

### *Vgll2* and *Tead1* mouse mutants display loss of Hcrt and Npvf expression

To probe the role of *Vgll2* and *Tead1* during hypothalamus development we knocked them out in the mouse. While constitutive *Vgll2* mutants survive until adulthood,^84^ constitutive *Tead1* mutants die at E12.5.^85^ Therefore, we knocked out *Vgll2* and *Tead1* conditionally in the CNS, by crossing floxed alleles generated by EUCOMM^86^ to *Sox1-Cre*; an early, pan-CNS deleter strain.^87^ Conditional *Vgll2* (*Vgll2^cKO^*) and *Tead1* (*Tead1^cKO^*) both survived postnatally. Neither *Vgll2^cKO^* nor *Tead1^cKO^*mutants displayed apparent gross brain anatomical abnormalities during developmental or early postnatal stages, but hydrocephaly was apparent in *Tead1^cKO^*at P20.

To uncover effects upon terminal differentiation we first analysed postnatal stage P20 by immunostaining. In *Vgll2^cKO^* we observed a 78% loss of Hcrt neurons and a complete loss of Npvf neurons, while *Tead1^cKO^*showed a complete loss of both Hcrt and Npvf neurons (Figure 2A-J, Table S1). To determine if Hcrt and Npvf neurons were generated during embryogenesis but lost, due to cell death or de-differentiation, we analysed P0 and E14.5–the earliest stage of robust Hcrt and Npvf immunostaining. At P0, the phenotypes were similar to P20, while at E14.5 the effects were even stronger, with a complete absence of both Hcrt and Npvf neurons in both mutants (Figure 2K-T, S6A-L).

**Figure 2.**
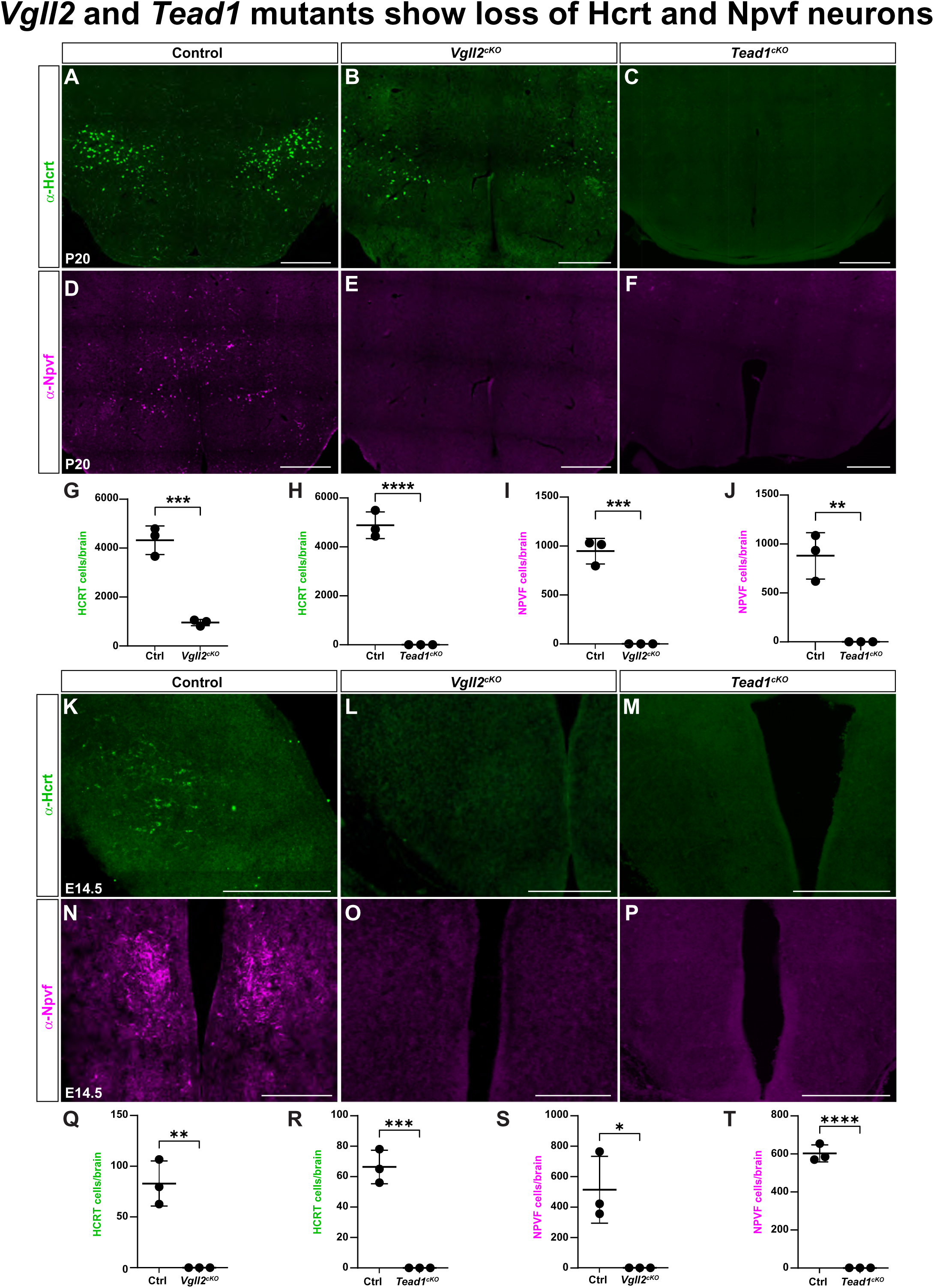
*Vgll2^cKO^*and *Tead1^cKO^* mouse mutants show loss of Hcrt and Npvf neurons. **(A-F)** Hcrt and Npvf expression in the P20 mouse hypothalamus in control, *Vgll2^cKO^* and *Tead1^cKO^*. (**G-J**) Quantification of Hcrt and Npvf neurons in the P20 mouse brain by serial sectioning, showing significant reduction of Hcrt and Npvf cells (Student’s two-tailed T-test; mean+/-SD; ** = p<0.01, *** =p≤0.001, **** = p≤0.0001; n = 3 brains analysed). **(K-P)** Hcrt and Npvf expression in the E14.5 mouse hypothalamus in control, *Vgll2^cKO^* and *Tead1^cKO^*. (**Q-T**) Quantification of Hcrt and Npvf neurons in the E14.5 mouse brain by serial sectioning, showing significant reduction of Hcrt and Npvf cells (Student’s two-tailed T-test; mean+/-SD; * = p<0.05, ** = p<0.01, *** =p≤0.001, **** = p≤0.0001; n = 3 brains analysed). Scale bars = 200μm.

Because of the well-known genetic and molecular interaction between VGLL and TEAD factors,^76,77^ and the overlap in expression and function between *Vgll2* and *Tead1* in the hypothalamus identified herein, we decided to probe for possible genetic interaction. To this end, we generated *Vgll2^cKO/+^;Tead1^cKO/+^*trans-heterozygotes. First, we analysed *Vgll2^cKO/+^* and *Tead1^cKO/+^* heterozygotes at P20 and observed that while *Vgll2^cKO/+^* heterozygotes did not display loss of Hcrt or Npvf, *Tead1^cKO/+^*heterozygotes showed a reduction of both neuronal cell types (Figure S7A-F). This effect was further enhanced in *Vgll2^cKO/+^;Tead1^cKO/+^* trans-heterozygotes (Figure S7A-F). These findings demonstrate that both *Vgll2* and *Tead1* are crucial for the generation of Hcrt and Npvf neurons and that they interact genetically regarding this function.

### sn-RNA-seq analysis points to a late role for *Vgll2* and *Tead1* in the pT1-1/EN21/Hcrt/Npvf pathway

To determine if the reduction of Hcrt and Npvf immunostaining reflected reduction of mRNA expression, and to probe the mutant phenotypes more comprehensively, we performed sn-RNA-seq analysis. Like the immunostaining, we first focused upon postnatal stages to probe the generation of most, if not all neuronal sub-types. We dissected the hypothalamus from control (*Vgll2^fl/+^* or *Tead1^fl/fl^*) and *Vgll2^cKO^*or *Tead1^cKO^* mutant animals, at P25 (*Vgll2^cKO^*) or P20 (*Tead1^cKO^*), with two animals for each genotype. Nuclei were sequenced on the 10xGenomics Chromium platform and after filtering 162,552 high-quality nuclei were obtained.

Analyses of the sn-RNA-seq data did not reveal any overt changes to the repertoire of broader cell types in either mutant (Figure 3A-B, S8A-B, Table S2). Differential expression gene (DEG) analysis revealed 324 DEGs, when comparing *Vgll2^cKO^* to control; 138 of which were upregulated and 186 downregulated (Table S3). In *Tead1^cKO^* animals we observed 92 DEGs; 39 upregulated and 53 downregulated (Table S3). *Hcrt* did not emerge as downregulated by DEG analysis in either mutant, and *Npvf* only in *Vgll2^cKO^*, likely due to the scarcity of these cells in the mature hypothalamus. However, violin plots revealed loss/reduction of *Hcrt* and *Npvf* nuclei in both mutants (Figure 3E-F). We did not observe altered expression of *Dbx1*, *Ebf2* and *Lhx9*, or any of the other proposed pT1-1/EN21 pathway TF genes in either mutant (Table S3).

**Figure 3.**
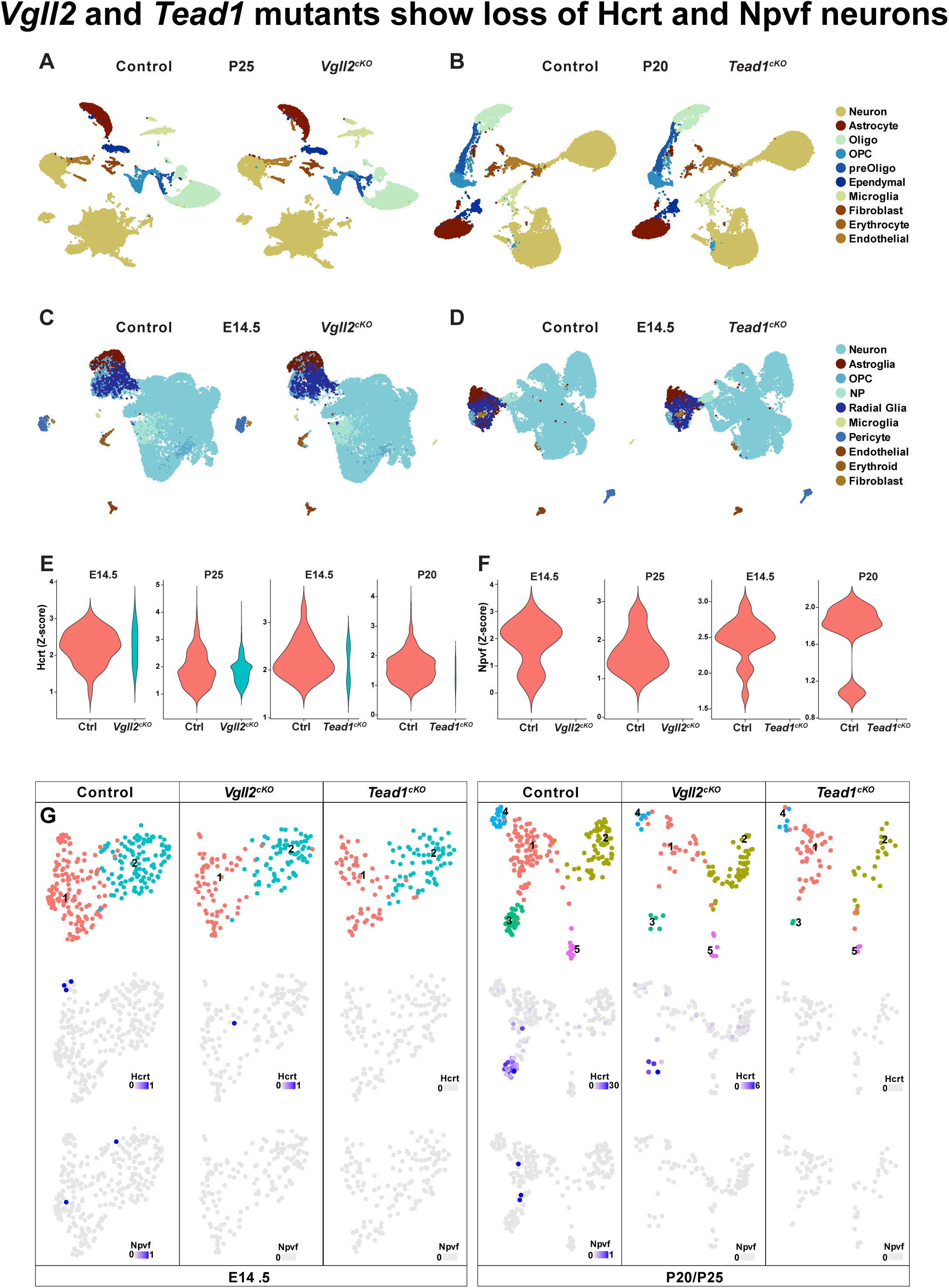
sn-RNA-seq analysis confirms the loss of Hcrt and Npvf neurons in *Vgll2^cKO^* and *Tead1^cKO^* mouse mutants. **(A-D)** sn-RNA-seq analysis of mouse hypothalamus at P25, P20 and E14.5, in control, *Vgll2^cKO^* and *Tead1^cKO^*. No apparent changes in major cell types are observed in either mutant at embryonic or postnatal stages (OPC, oligodendrocyte precursor cells; NP, neural precursor). **(E-F)** Violin plots show reduction/loss of *Hcrt* **(E)** and *Npvf* **(F)** nuclei in both *Vgll2^cKO^* and *Tead1^cKO^* mutant mice, at both embryonic and postnatal stages. **(G)** Analysis of the EN21 neuronal cluster in *Vgll2^cKO^*and *Tead1^cKO^* mutant mice at E14.5 and P20/P25 revealed that most EN21 cluster nuclei appear to be generated also in the mutants.

To better understand the origins of the loss/reduction of *Hcrt* and *Npvf* neurons we conducted sn-RNA-seq analysis at E14.5. We dissected the hypothalamus from two control (*Vgll2^fl/+^* or *Tead1^fl/fl^*) and two *Vgll2^cKO^* or *Tead1^cKO^* mutant animals, at E14.5, and obtained a total of 73,931 high-quality nuclei. Like P20/P25, we did not find overt changes to the repertoire of broader cell types (Figure 3C-D, S8A-B, Table S2). DEG analysis comparing *Vgll2^cKO^*to control revealed 173 DEGs; 103 upregulated and 70 downregulated, which included *Npvf* and *Hcrt* (Table S3). In *Tead1^cKO^* animals we observed 194 DEGs; 59 upregulated and 135 downregulated, which again included *Npvf* and *Hcrt* (Table S3). Violin plots also revealed loss/reduction of *Hcrt* and *Npvf* nuclei in both mutants (Figure 3E-F). Notably, three other neuropeptide genes, *Qrfp*, *Avp* and *Pmch*, were also downregulated in *Tead1^cKO^* (Table S3). Like the P20/P25 results, we did not observe altered expression of *Dbx1*, *Ebf2* and *Lhx9*, or any of the other proposed pT1-1/EN21 pathway genes in either mutant (Table S3).

The loss/reduction of *Hcrt* and *Npvf* expressing nuclei in both mutants raised the question of the underpinnings of these phenotypes. Because two of the key marker genes defining the EN21 cluster, *Lhx9* and *Rfx4*,^30^ were not downregulated in either mutant, we sought to probe the development of the EN21 cluster in the mutants. As previously described^30^, we identified a small subset of EN21 nuclei (*Lhx9^+^/Rfx4^+^*) within the hypothalamus. At E14.5, the EN21 nuclei could be sub-divided into two Leiden clusters (Figure 3G). Both the *Vgll2^cKO^* and *Tead1^cKO^* mutant samples contained EN21 nuclei, which clustered like control across the two Leiden clusters (Figure 3G; note that for control, all four samples were combined, and hence more control EN21 nuclei are observed). Expression of *Hcrt* and *Npvf* was observed in subsets of control EN21 nuclei, reduced in *Vgll2^cKO^* and absent in *Tead1^cKO^*mutants (Figure 3G). Analysing the P20/P25 samples revealed a similar picture, with a small subset of EN21 nuclei present in the control samples, albeit with a more complex clustering into five Leiden clusters (Figure 3G). Both the *Vgll2^cKO^*and *Tead1^cKO^* mutant samples contained EN21 nuclei, which clustered like control across the five Leiden clusters (Figure 3G). Expression of *Hcrt* and *Npvf* was again observed in subsets of control nuclei, with *Hcrt* reduced and *Npvf* lost in *Vgll2^cKO^*, and both neuropeptide genes absent in *Tead1^cKO^* mutants (Figure 3G).

The sn-RNA-seq results generally align with the immunostaining results, showing loss/reduction of *Hcrt* and *Npvf* expression in both mutants. The one marked difference was that while Hcrt immunostaining was completely absent at all stages in *Tead1^cKO^* mutants, the sn-RNA-seq did uncover a small number of *Hcrt* mRNA expressing cells. The sn-RNA-seq analysis indicates restricted but key roles for *Vgll2* and *Tead1* late in the proposed pT1-1/EN21 genetic pathway, acting not to generate the EN21 cluster but rather to govern Hcrt and Npvf neuronal cell fates.

### *VGLL2*-*TEAD1* co-misexpression induced EN21/HCRT neurons in human hypothalamic organoids

Next, we decided to start addressing the evolutionary conservation of the proposed pT1-1/EN21/Hcrt/Npvf pathway into humans and the potency of *VGLL2* and *TEAD1* in driving human HCRT and NPVF neuron specification. To this end, we developed a hypothalamic organoid system (HypoOrgs), derived from human induced pluripotent stem cells (hiPSCs). We deployed a stepwise differentiation protocol,^70^ based upon dual-SMAD and WNT inhibition, to trigger neural induction and rostral CNS development, followed by SHH activation, to guide ventral CNS development. These initial patterning steps were followed by stepwise maturation cues (Figure 4A). At Day 60 (D60), immunostaining and sn-RNA-seq analyses revealed that most major hypothalamic cell types were generated in the HypoOrgs (Figure 4B-I, S9A-B). In line with their *in vivo* expression profiles, expression of TEAD1 was broad whereas VGLL2 expression was quite restricted (Figure 4D-E, 4H). HCRT expression was quite limited, and only detected by sn-RNA-seq, while NPVF expression was not observed at all (Figure 5A, 5I).

**Figure 4.**
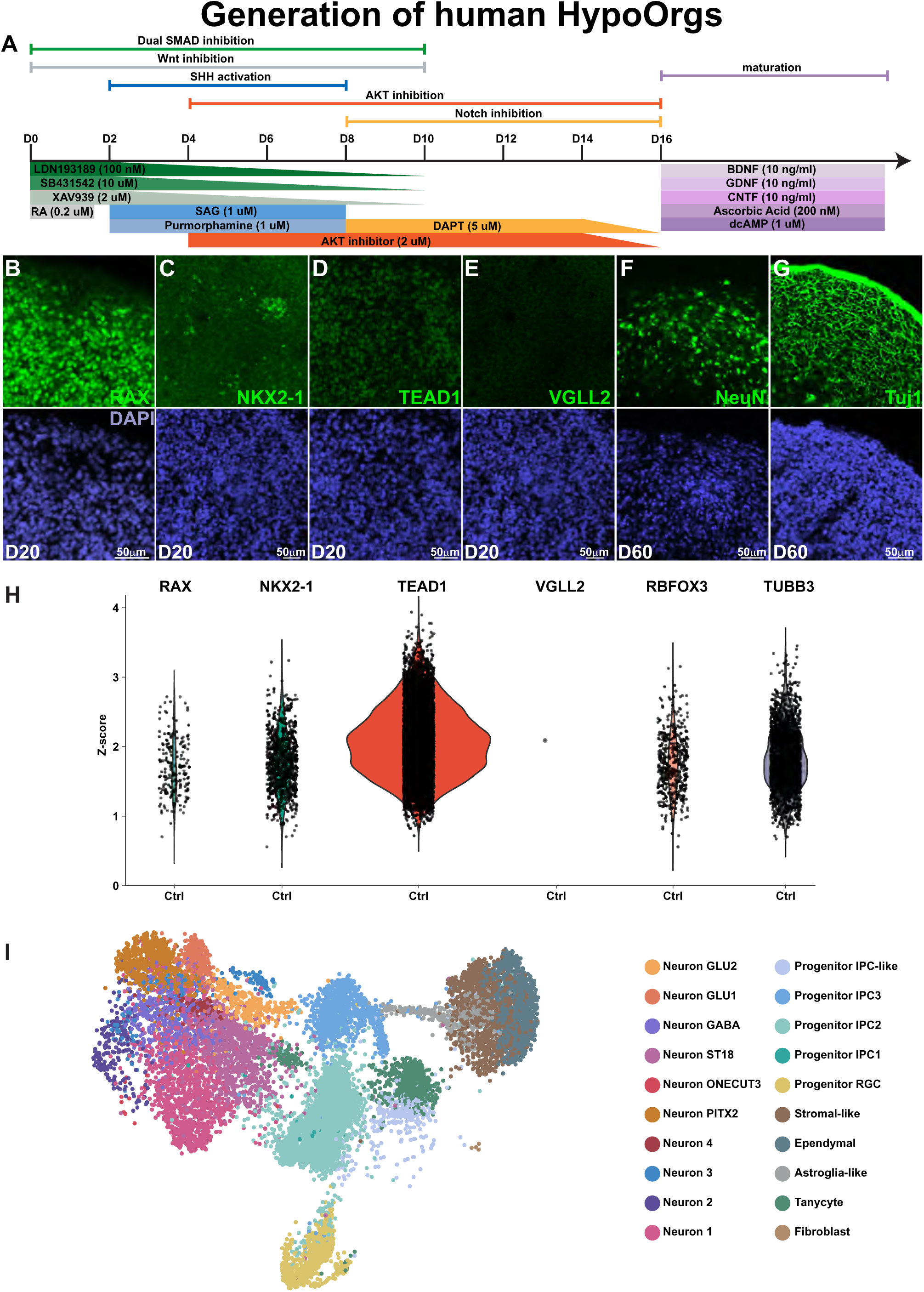
Generation of human hypothalamic organoids. **(A)** Differentiation protocol used to drive hiPSCs to hypothalamic organoids (HypoOrgs). **(B-G)** Marker analysis by immunostaining. **(H)** sn-RNA-seq analysis reveals major cell type generation in HypoOrgs at D60. **(I)** UMAP showing the presence of major cell types in the control HypoOrgs.

**Figure 5.**
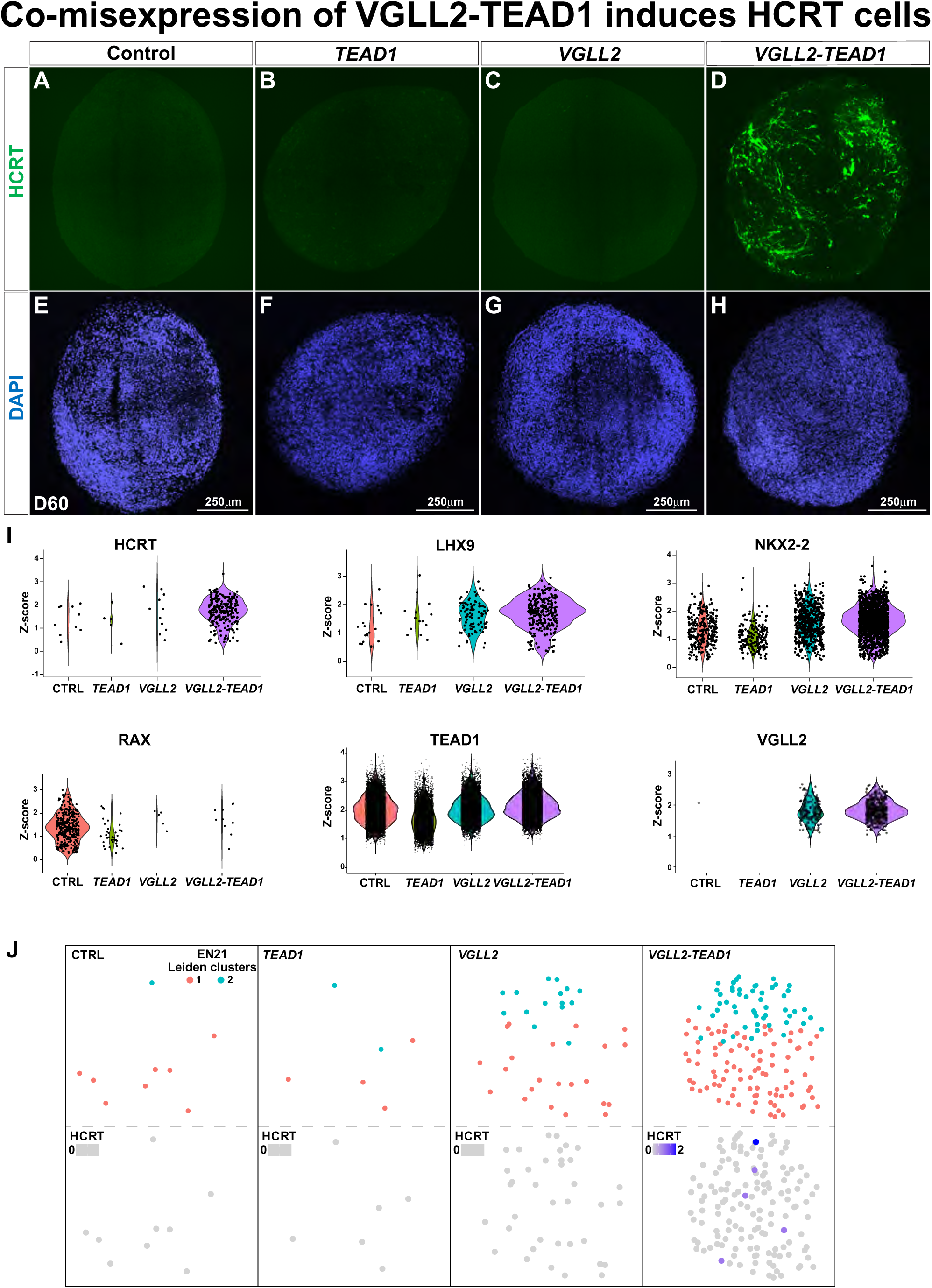
*VGLL2-TEAD1* co-misexpression in human hypothalamic organoids induces EN21/HCRT neuron differentiation. **(A-H)** Staining for HCRT and DAPI in 20μm sections of HypoOrgs at D60. **(I)** Violin plots based on sn-RNA-seq analysis of D60 HypoOrgs, showing nuclei expressing *HCRT*, *LHX9*, *NKX2-2*, *RAX*, *TEAD1* or *VGLL2*. **(J)** EN21 cluster nuclei (*LHX9*^+^/*RFX4*^+^) in control and misexpressing HypoOrgs. *VGLL2* single and *VGLL2*-*TEAD2* co-misexpression induces extra EN21 neurons, a subset of which express *HCRT*.

Next, we generated lentiviruses expressing *VGLL2* or *TEAD1* under control of the doxycycline (dox) inducible TRE promoter^88^ (Info S1). Infection of a previously generated hiPSC line expressing the TRE activator rtTA constitutively from the AAVS1 locus^89^ allowed for dox-inducible VGLL2 or TEAD1 expression (Figure S10A-F’’). The lentivirus vectors also contained EGFP (in the VGLL2 vector) or mCherry (TEAD1) marker genes, constitutively expressed, allowing for the generation of single and double expressing hiPSC lines by FACS sorting. Due to silencing, we found that two rounds of FACS sorting were required to obtain lines that expressed VGLL2 and TEAD1 robustly in most cells (Figure S10G-I).

To test the sufficiency of *VGLL2* and *TEAD1* during HypoOrg differentiation, we induced expression of these genes at D4 or D10, and monitored the effects at D60, using immunostaining and sn-RNA-seq. As outlined above, at D60 in control HypoOrgs, HCRT expression was quite limited and NPVF not detected, by immunostaining or sn-RNA-seq (Figure 5A, 5I). As anticipated from the highly limited endogenous expression of *VGLL2* in HypoOrgs and the lack of evidence for activation of *Vgll2* by *Tead1* in mouse, *TEAD1* misexpression did not induce *HCRT* or *NPVF* cells (Figure 5B). Somewhat surprisingly, given the endogenous expression of *TEAD1* in control HypoOrgs, single *VGLL2* misexpression did also not induce *HCRT* or *NPVF* expression (Figure 5C). However, *VGLL2-TEAD1* co-misexpression induced expression of *HCRT*, evident by immunostaining, with apparent axon/dendrite extensions (Figure 5D). *NPVF* was not induced even in *VGLL2-TEAD1* co-misexpressing HypoOrgs (Table S5). The induced HCRT (iHCRT) neurons were apparent at earlier stages, D40, but displayed shorter projections, and maintained at later stages, D80 (Figure S11A-P). sn-RNA-seq confirmed the immunostaining results, showing induction of HCRT expressing nuclei (Figure 5I).

Focusing further on the sn-RNA-seq data for control and *VGLL2-TEAD1* co-misexpressing HypoOrgs, DEG analysis identified 551 DEGs, with 217 upregulated, including *HCRT*, and 334 downregulated (Table S5). As anticipated, *VGLL2* was DE in *VGLL2* and *VGLL2-TEAD1* double misexpression (Table S5), although violin plots reveal quite limited numbers of nuclei showing *VGLL2* expression in either genotype (Figure 5I). These findings contrast with the robust expression of VGLL2 in the lentivirus infected and double FACS sorted hiPSCs (Figure S10G-H), pointing to substantial silencing during the 60-day long differentiation protocol. *TEAD1* was not DE in *TEAD1* or *VGLL2-TEAD1* HypoOrgs, and not apparently elevated in violin plots, likely due to the high endogenous levels already present in the HypoOrgs (Figure 5I, Table S5).

By contrast to the mouse mutant analysis, which did not reveal significant changes in any of the proposed pT1-1/EN21 pathway genes, DEG analysis revealed that both the *VGLL2* single and *VGLL2-TEAD1* co-misexpression resulted in upregulation of *LHX9*, *HMX2/3* and *NKX2-2*, and downregulation of *RAX* and *DBX1* (Figure 5I, Table S5). The regulation of some of the pT1-1/EN21 pathway genes prompted us to analyse the EN21 cluster in the HypoOrgs. In control HypoOrgs, we observed a small set of EN21 nuclei, which was not markedly altered in *TEAD1* misexpression (Figure 5J). By contrast, the *VGLL2* single and in particular *VGLL2-TEAD1* co-misexpression resulted in a marked increase in EN21 nuclei, a subset of which also expressed *HCRT* (Figure 5J).

Previous studies have indicated several different subtypes of Hcrt neurons, evident by different electrical and pharmacological signatures, transcriptional profiles, and axonal projections.^90–93^ However, attempts to identify Hcrt subclusters based upon mouse sc-RNA-seq data did not clearly reveal this diversity, likely due to the small number of Hcrt cells available for analysis and confounding effects of sex differences in the samples.^14^ With much greater numbers of sn/sc-RNA-seq data now publicly available, we decided to probe the diversity of human HCRT neurons and compare the iHCRT neurons to their *in vivo* counterparts. To this end, we downloaded published human hypothalamus sn/sc-RNA-seq data sets,^26,29^ holding some 588,552 cells/nuclei in total: 155,183 embryonic and 433,369 adult. Because the hiPSC line used in our study was male we only analysed male cells. We extracted HCRT expressing male cells, totalling 2,323 cells/nuclei: 1,637 adult and 686 embryonic. In line with the notion of HCRT subtype diversity, our meta-analysis revealed that the adult human HCRT neurons dispersed into 12 distinct UMAP clusters, a subset of which also contained embryonic HCRT nuclei (Figure 6A-D, 6F). Gene ontology (GO) analysis of the DEGs defining the 12 different clusters revealed GO term enrichment for Biological Processes, Cellular Components and Molecular Processes for most clusters. Strikingly, in several cases the enriched GO terms involved axon pathfinding, synapse formation and synaptic transmission (Figures S12-14, Table S6). HypoOrg iHCRT nuclei contributed to many, but not all the adult clusters and were particulary enriched bot clusters #2 and 5 (Figure 6B-D). A subset of DEGs triggered by *VGLL2-TEAD1* co-misexpression overlapped with HCRT cluster marker genes, and in some cases were up- or downregulated in correlation to the cluster profile triggered by *VGLL2-TEAD1* co-misexpression e.g., up for cluster #2 and down for cluster #10 (Figure 6E).

**Figure 6.**
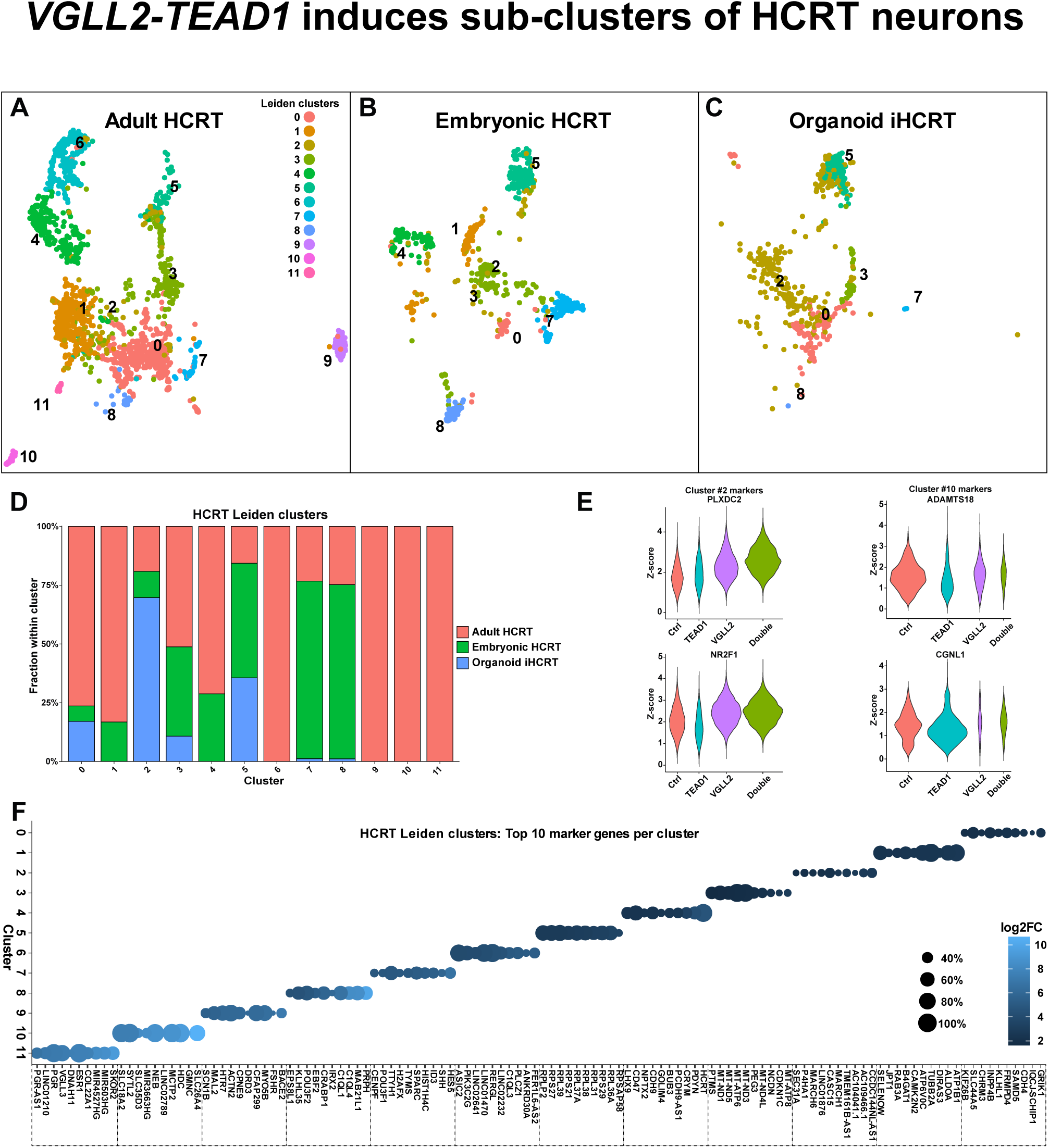
*VGLL2-TEAD1* co-misexpression induces HCRT sub-cluster neuron differentiation. **(A-C)** UMAP analysis comparing human adult, embryonic and HypoOrg *HCRT* nuclei. **(A)** Human adult *HCRT* nuclei cluster into 12 Leiden clusters. **(B)** Embryonic *HCRT* and HypoOrg *iHCRT* **(C)** nuclei contribute to many of the adult clusters, but show limited, if any contribution to some clusters. **(D)** Proportions of adult, embryonic and HypoOrg HCRT nuclei of each cluster. **(E)** Violin plots of HCRT cluster #2 and #10 marker genes in the HypoOrgs of different genotypes. **(F)** Dot plot of top 10 marker genes for each of the 12 adult HCRT clusters.

These results revealed that *VGLL2* and *TEAD1* could act in a combinatorial code in human HypoOrgs to trigger the generation of EN21 and iHCRT neurons, and that these iHCRT neurons displayed transcriptional profiles that overlapped with many, but not all putative subtypes of adult human HCRT neurons.

## DISCUSSION

### The pT1-1 progenitor domain

Previous studies, using *in situ* hybridization, identified the region generating mouse Hcrt neurons as located in the dorsal-most *Nkx2-1* region in the developing tuberal hypothalamus, in an anterior-posterior stripe of *Lhx9* expression and dorsally bordered by *Lhx6*, *Lhx8* and *Arx*.^18^ Building upon this mapping we analysed gene expression in the tuberal hypothalamus, using a number of different gene expression data sets and immunostaining, and integrated *Vgll2* and *Tead1* into a spatio-temporal framework for dorso-tuberal development. Building upon recent progenitor definitions, which identified nine progenitor domains in the developing hypothalamus, including two for the tuberal hypothalamus, pT1 and pT2, we propose a sub-division of the pT1 domain into pT1-1 (dorsal-most) and pT1-2 (intermediate), with pT1-1 being *Nkx2-2*^+^/*Six6*^-^ and pT1-2 being *Nkx2-2*^-^/*Six6*^+^/*Tbx3*^-^, and with pT2 ventral-most and *Tbx3*^+^ (Figure 1M). Given the immense cellular complexity of the hypothalamus, with hundreds of transcriptionally distinct differentiated cell subtypes, further sub-division of the nine progenitor domains is arguably anticipated. We propose that the pT1-1 progenitor domain generates the EN21/Hcrt/Npvf glutamatergic neuron cluster, which contributes neurons into the two mature dorso-tuberal nuclei, the DMH and LH. Baring *in vivo* lineage analysis we can only speculate about the pT1-1 lineage tree, whether it deploys indirect and/or direct neurogenesis, its lineage size, and the lineage relationship between Npvf and Hcrt neurons.

### The emerging pT1-1/EN21/Hcrt/Npvf genetic pathway

We propose a genetic pathway from pT1-1 progenitors to EN21/Hcrt/Npvf neurons, from broader hypothalamic patterning genes expressed in progenitors i.e., *Rax*, *Dbx1*, *Nkx2-1* and *Nkx2-2*, transitioning to more restricted and chiefly postmitotically expressed genes i.e., *Hmx2/3*, *Lhx9* and *Vgll2*. In addition, two genes initially expressed broadly in progenitors, *Tead1* and *Rfx4*, display restricted postmitotic expression in the EN21 cluster. Apart from *Rax* and *Dbx1*, these eight TFs continue expressing in the EN21 cluster into adults. The functional importance of this proposed genetic pathway is supported by the loss of mouse Hcrt neurons in mutants for four of these eight TF genes: *Dbx1*, *Lhx9, Tead1* and *Vgll2*. How this proposed genetic pathway is connected remains to be determined, but evidence points to *Dbx1* regulating *Lhx9*,^73^ and *Vgll2-Tead1* regulating *Nkx2-2*, *Lhx9* and *Hmx2/3*, at least as evidenced from our co-misexpression results in HypoOrgs. Moreover, while the mouse *Tead1* and *Vgll2* mutant analysis did not reveal an involvement of these two genes in the specification of the broader EN21 neuronal cluster, the misexpression in HypoOrgs indicated involvement herein.

The *Ebf2* gene was also found to be important for Hcrt neuron specification,^71^ but our expression analysis did not shed light on its place within the cascade. The spatial RNA-seq data shows *Ebf2* to be expressed in subsets of cells in many hypothalamic regions, both in progenitor and differentiating regions. The sc/sn-RNA-seq data show that *Ebf2* is transiently expressed in subsets of progenitors and intermediate progenitors, with some transient overlap with *Lhx9* in a subset of intermediate progenitors. Given the loss of Hcrt neurons in *Ebf2* mutants we envision that it may be transiently expressed in intermediate progenitors in the pT1-1/EN21 pathway.

*Hmx2* and *Hmx3* appear to be genetically redundant in the CNS and have been found to play important roles in developing hypothalamus.^94^ Their expression in the tuberal hypothalamus is different along the anterior-posterior axis, traversing the entire dorso-ventral region posteriorly, while being restricted to the mid- and dorso-tuberal region anteriorly (Figure S2A). In line with this expression pattern, *Hmx2/3* were found to be important for the specification of neuropeptide neurons in the ventro-tuberal region, in the Arcuate nucleus.^94^ The possible role of *Hmx2/3* in the dorso-tuberal region has not been addressed. *Rfx4* is initially expressed broadly in the CNS, in progenitor regions, and mutants display DV patterning defects.^95–97^ However, *Rfx4* becomes increasingly restricted during later stages and in the hypothalamus is observed in a subset of neuronal clusters, being a marker for the developing EN21 cluster,^30^ as well as the related C66-18:Rfx4.GLU-4 cluster, identified in the adult mouse.^5^ The role of mouse *Rfx4* during these later stages has not been addressed, but interestingly, GWAS analysis of ∼58,000 individuals scoring for human chronotypes mapped seven out of the 15 SNPs identified to the first intron of *RFX4*.^98^

Previous studies indicated the presence of different subtypes of Hcrt neurons, evident by different electrical and pharmacological signatures, transcriptional profiles, and axonal projections.^90–93^ This notion is supported by our meta-analysis of human sn/sc-RNA-seq data, indicating several transcriptionally distinct sub-clusters of HCRT neurons. It is possible that the genes in the pT1-1/EN21/Hcrt/Npvf pathway proposed herein, perhaps in different combinations of e.g., splice forms, protein levels and/or protein modifications, may be sufficient to diversify HCRT neurons. However, it is also possible, arguably probable, that additional regulators are involved in the diversification of HCRT subtypes.

*Vgll2* was previously identified as expressed also in the mid-tuberal region, subsequently forming the VMH.^12^ Like in the dorso-tuberal region, *Vgll2* expression in the developing VMH appears chiefly postmitotic, and is evident within the recently proposed EN7 and EN8 glutamatergic clusters.^30^ We did not investigate the potential role for *Vgll2* in the VMH to any length, but note that none of the three neuropeptide genes *Adcyap1*, *Pdyn* and *Tac1*, which are strongly expressed in the EN7 and/or EN8 clusters,^30^ were significantly affected in *Vgll2^cKO^*mutants, as detected by sn-RNA-seq (Table S3).

### The EN21/HCRT/NPVF lineage in humans and in HypoOrgs

In humans, like mouse, immunostaining has identified HCRT and NPVF neurons in the adult hypothalamus, in the DMH and LH regions.^60,61,79–81^ While detailed spatial gene expression analysis of human embryos is still emerging, sn/sc-RNA-seq data supports a similar developmental and genetic trajectory of a human pT1-1/EN21/HCRT/NPVF domain as of that proposed herein for mouse. However, as anticipated, there are major differences between mouse and human regarding the number of Hcrt neurons, with Hcrt numbers sitting at ∼4,000 in the adult mouse hypothalamus and at ∼70,000 in the adult human hypothalamus.^61^ How this scaling is achieved is difficult to predict, and similar in challenge to all mouse-human comparisons, and it may involve both increased numbers of pT1-1 progenitors and larger lineages.

To determine the role of the pT1-1/EN21/HCRT/NPVF pathway genes in humans and to generate iHCRT neurons in large numbers, hiPSCs provide a potential model system. Many studies have driven hiPSCs into hypothalamic differentiation *in vitro*, both in 2D and 3D cultures,^69^ referred to as HypoOrgs herein. These studies have deployed sequential patterning cues, attempting to mimic the normal development of the mammalian hypothalamus, and have demonstrated expression of hypothalamic patterning and cell-specific markers.^69^ These include many neuropeptide genes, including HCRT, although as anticipated HCRT cells were observed in small percentages.^70^ Only a few studies have attempted to generate specific hypothalamic nuclei,^99^ and in general the complex multi-step patterning of the embryonic hypothalamus^2,3,31,39–48^ makes it an arguably particularly challenging tissue to model in organoids. Systematic analysis of different culture conditions by comprehensive sn/sc-RNA-seq analysis may help progress the field further and enable more efficient induction of HCRT and other cell subtypes. However, recent comprehensive combinatorial morphogen culturing of CNS organoids indicates that morphogens alone are often insufficient in generating only one neural cell type,^100^ pointing to the requirement of TF guidance for specific cell type generation.

To test the roles of *VGLL2* and *TEAD1* we misexpressed them in human HypoOrgs. As anticipated from the limited expression of *VGLL2* in control HypoOrgs, single misexpression of *TEAD1* did not induce iHCRT cells. Surprisingly, given the endogenous expression of *TEAD1* mRNA and protein in HypoOrgs, we found that *VGLL2* misexpression was also insufficient to induce iHCRT cells. This may be due to that the endogenous TEAD1 protein levels are too low and/or that the wrong mRNA splice variant is expressed. However, *VGLL2-TEAD1* co-misexpression, delivered from recombinant lentiviruses, could trigger iHCRT production. We observed a low proportion of iHCRT cells in *VGLL2-TEAD1* co-misexpressing HypoOrgs, likely due to silencing of the lentiviruses, a well-recognized challenge in extended organoid cultures.^101^ Indeed, while we observed broad and robust VGLL2 and TEAD1 expression in the lentivirus infected, double FACS-sorted hiPSC lines at the iPSC culturing stage, expression at D60 was limited. Development of more robust transgenic systems will likely increase the number of iHCRT neurons generated. Moreover, both the mouse and human gene expression analysis revealed that *TEAD1* is expressed before *VGLL2*, first in progenitors and then in differentiating neurons, a sequence that may be important and that is challenging to precisely re-create with current technologies.

We did not observe any NPVF cells in our HypoOrgs, even in the co-misexpression scenario. No other HypoOrg studies have specifically analysed NPVF expression by immunostaining, so it is unclear if this is a common limitation. The reason for the lack of NPVF cells may relate to timing, since in contrast to mouse, where Npvf and Hcrt appear in the same developmental time-window, in humans, the sn/sc-RNA-seq data indicates that *NPVF* onset occurs later than *HCRT*, at PCW10 as opposed to PCW6. However, even at D80 in HypoOrgs we could still not detect NPVF. Alternatively, NPVF expression may depend upon target-derived signals from other parts of the CNS, like what has been observed for the FMRFa neuropeptide in *Drosophila*,^102^ and/or circulating hormones from the periphery, which may be missing from the culture medium.

### iHCRT neurons for human physiology studies and cell therapy

There are currently 115 ongoing regulatory approved clinical trials involving hiPSC cell therapy, 20 out of which involve cell transplantation into the human CNS.^103^ While these studies are still ongoing, recent updates on the progress of e.g., hiPSC-derived dopamine cells into Parkinson’s Disease (PD) patients are encouraging.^104,105^ However, the case of cell transplantation into PD illustrates the challenges emerging from the cellular complexity in the CNS. Specifically, several different subtypes of midbrain dopaminergic neurons have been identified *in vivo*, with some subtypes apparently more affected than others in PD.^106^ It is unclear if this *in vivo* and disease complexity will constitute a future hurdle in cell therapy treatments.

Similar issues may be important to consider regarding cell therapy for narcolepsy. Previous studies have pointed to different HCRT subtypes,^90–93^ and our sn/sc-RNA-seq meta-analysis herein identified 12 transcriptionally distinct HCRT sub-clusters in humans. These findings indicate that the different Hcrt neuron subtypes evident from previous functional studies are underpinned by distinct gene expression profiles. We speculate that one main driver of this sub-clustering may relate to axon/dendrite projections, as recent studies in mouse reveal that different subsets of Hcrt neurons project to different CNS regions, identifying four major projection subtypes.^92^ Indeed, our GO analysis of the DEGs defining the 12 different HCRT cell clusters revealed many GO terms involving axon pathfinding and synapse formation, hence lending some support for this notion. We find that *VGLL2-TEAD1* co-misexpression can generate iHCRT neurons in HypoOrgs, but that these do not overlap fully with all the *in vivo* sub-clusters. However, how/if the sn/sc-RNA-seq clusters directly correspond to different *in vivo* defined subtypes, as defined by e.g., pharmacology, electrophysiology and connectome, is currently difficult to assess.

HCRT immunostaining and *in situ* hybridisation studies of narcoleptic individuals have indicated that symptoms begin to occur at an extensive, but partial, loss of HCRT neurons.^90^ However, it is not clear if the HCRT cells lost belong to a particular subtype(s). Previous studies involved transplanting Hcrt-GFP cells from one mouse strain into Hcrt cell-ablated mouse hosts.^66,67^ These cell injections were made into one of the key target areas of Hcrt projections, the Dorsal Raphe nucleus, and resulted in some alleviation of narcoleptic symptoms, specifically cataplexy. However, these studies presumably involved mixtures of Hcrt subtypes. The combination of further analysis of human HCRT subtypes, including the potential selective subtype loss in narcoleptic patients, improved methods for generating iHCRT neurons, a deeper understanding of the physiology of human HCRT neurons, and comprehensive testing of cell injection target regions will likely all be necessary for the development of cell therapy for narcolepsy.

## Acknowledgements

We are grateful to UC Davis KOMP and RIKEN mouse stock centres, as well as Ernst Wolvetang for sharing reagents and advice. We are grateful to Qing-Feng Wu for advice on gene expression analysis. We are grateful to Jiaojiao Liu, Kirra Csurhes and Lili Wang for early aspects of the projects. Helen Ekman and Angelika Christ-Velde provided excellent technical assistance. We thank Qing-Feng Wu and Massimo Hilliard for critically reading the manuscript.

## Author contributions

Conceptualization: ST. Software: RW, AS and BB. Methodology: RW, AS, AG, CI-G, BB and ST. Formal analysis: RW, AS, BB and ST. Investigation: RW, AS, ES, BB and ST. Resources: MP, MB and ST. Data Curation: RW, AS and ST. Writing – original draft preparation: RW, AS and ST. Writing – review and editing: RW, AS, ES, BB, AG, CG-I, MB, MP and ST. Visualization: RW and ST. Supervision: ST, MP and MB. Funding acquisition: MP, MB and ST.

## METHODS

### Mouse stocks

The *Tead1^fl/fl^* and *Vgll2^fl/fl^* alleles (*Tead1^tm1a(KOMP)Wtsi^* and *Vgll2^tm1a(KOMP)Wtsi^*) were generated by the Knockout Mouse Project (KOMP) consortium ^107^. Embryonic stem cells blastocyst injections, chimeric and germline mice generation was conducted by the UC Davis KOMP Repository (UC Davis, USA) for *Tead1* and *Vgll2*. The *lacZ-neo* cassette in *Tead1* was removed by Flp mediated deletion at UC Davis KOMP Repository. The *Sox1-Cre* line^87^ was provided by RIKEN (RBRC05065). We observed deletion of the targeted exons by PCR of genomic DNA and/or loss of Tead1 and Vgll2 immunostaining in the hypothalamus in the mutants. Mice were maintained at The University of Queensland, Queensland Brain Institute animal facility. All mouse procedures were approved by the regional animal ethics committee (AEC approval number QBI/191/20). Detection of the vaginal plug in the morning was set as 0.5 days post coitum i.e., E0.5. Primers for genotyping were: Cre1: GCG GTC TGG CAG TAA AAA CTA TC; Cre2: GTG AAA CAG CAT TGC TGT CAC TT; Tead1: GGG ACG TGC TGA CAT TTT CT; Tead2: CTT GGG TGG TTT GGC TAA GA; Vgll2-3: GAG ATG GCG CAA CGC AAT TAA TG; Vgll2-4: GAG GAA AAC CAA ACC TTA CTG TAA GGA C; Vgll2-5: TGG CAA AAG CCT GGC TAG TTT TAG G; Vgll2-6: TCA TGT TTT CTG TGA TAC TTC ATT CCT G.

### Immunohistochemistry

Postnatal mice (P0 and P20) were perfused trans-cardially with ice-cold phosphate-buffered saline (PBS, pH 7.4), followed by ice-cold 4% paraformaldehyde (PFA) in PBS (pH 7.4). The brains were removed from the skull and transferred to a 15% sucrose solution in PBS at 4°C overnight. The following day, the brains were transferred to a 30% sucrose solution in PBS at 4°C and incubated until fully saturated, as indicated by sinking to the bottom. Once saturated, the brains were embedded in OCT Tissue-Tek (Sakura Finetek, Alphen aan den Rijn, Netherlands) and frozen in iso-pentane (prechilled with dry ice) and stored at -80°C.

For embryonic mice, heads were decapitated, and skulls were slightly cut opened before putting in 4% PFA. To enhance PFA penetration, fixation was assisted using the Pelco Biowave Microwave System. Following Biowave treatment using a standard fixation protocol #1, 150W microwave power in a pulsed, on-off-on cycle: 40 seconds on, 40 seconds off, and then another 40 seconds on). After fixation, heads were incubated in 4% PFA overnight at 4°C. The next day, heads were transferred to a 15% sucrose solution (in PBS) at 4°C overnight, followed by incubation in a 30% sucrose solution (in PBS) at 4°C until fully saturated. The samples were then embedded in OCT Tissue-Tek and frozen in iso-pentane (prechilled with dry ice) and stored at -80°C. All brains were sectioned at 20 µm thickness using a cryostat.

For organoids, on day of collection, samples were harvested by first removing the cultural medium and washed 3 times with PBS (pH 7.4) for 5 mins per wash. Organoids were then fixed in 4% PFA in PBS (pH 7.4) for 30 minutes at room temperature, washed twice with PBS, and were transferred to a 15% sucrose solution in PBS at 4°C overnight, followed by incubation in 30% sucrose solution at 4°C until fully saturated (sinking). Once saturated, organoids were embedded in OCT Tissue-Tek and frozen in iso-pentane (prechilled with dry ice) and stored at -80°C. All brains and organoids were sectioned at 20 µm thickness using a cryostat.

Prior to staining, cryosections were retrieved from -80°C and allowed to dry at room temperature for 30 mins. The slides were then placed in a 50°C oven for 2 hours to ensure proper adhesion. For immunostaining, sections underwent two rounds of photobleaching, where slides were covered with a thin layer of PBS, placed in closed tray and exposed to high-powered white LEDs for 30 mins. Slides were then treated with citrate buffer (10 mM, pH 6) at 90°C for 10 mins, followed by cooling at room temperature for 1 hour. Sections were then washed three times with PBS (5 mins each). For blocking, sections were incubated with blocking buffer (3% donkey serum in 0.2% Triton X-100 PBS) for 1 hour at room temperature (RT). Sections were incubated with primary antibodies diluted in blocking buffer overnight at RT. Fluorophore-conjugated secondary antibodies (FITC, Rhodamine-RedX, Cy5; Jackson ImmunoResearch, West Grove, PA, USA) were applied at 1:1,000 in blocking buffer for 3 hrs at RT and washed three times with PBS (5 mins each). For reducing autofluorescence, sections were treated with 0.5% Sudan Black B solution (in 70% ethanol) for 30 mins, followed by two washes with 70% ethanol (5 mins each). Counter nuclear staining was performed with Hoechst 33342 (Thermo Fisher Scientific #62249, 20mM, 1:1,000 dilution in PBS) for 10 mins then washed three times with PBS (5 mins each). Finally, slides were mounted in Eukitt Quick-hardening mounting medium (Sigma-Aldrich #03989) and stored at +4°C in the dark until microscopy imaging.

### Antibodies

Primary antibodies for mouse brain slices: rabbit anti-Tead1 (1:1,000, ab133533, Abcam), goat anti-Sox2 (Y17) (1:500, SC17320, Santa Cruz), rabbit anti-Orexin A (1:1,000, ab6214, Abcam), rabbit anti-Orexin (D6G9T) (1:500, #16743, Cell Signaling Technology), IgY anti-Hcrt (1:500, this study), rabbit anti-Vgll2 (1:500, PA5100693, Invitrogen), guinea pig anti-Vgll2 (1:250, this study), guinea pig anti-Npvf (1:500, this study), goat anti-Npvf (1:500, this study).

Primary antibodies for iPSCs/organoids: mouse anti-TTF1/Nkx2-1 (1:500, ab212888, Abcam), guinea pig anti-Rax (1:500, M229, TaKara), rabbit anti-Sox2 (1:500, ab92494, Abcam), mouse anti-βIII Tubulin (1:500, MAB1195, R&D systems), rabbit anti-Neun (D4G4O) (1:500, #24307, Cell Signaling Technology), chicken anti-mCherry (1:500, ab205402, Abcam), goat anti-GFP (1:500, ab6673, Abcam).

Antibodies to mouse Npvf and Vgll2 were generated by inserting DNA fragments encoding the full-length pre-pro-Npvf or Vgll2 proteins (both genes are predicted to only generate one protein variant; https://genome.ucsc.edu) into pGEX-4T3 and expressed in *E. coli* bacteria. ORFs were codon optimized for *E. coli* expression and synthesized by de-novo gene synthesis (Genscript, Piscataway, NJ, USA) (sequences in Suppl Info 1). SDS-PAGE-gel purified Gst-fusion proteins were injected into guinea pigs, rabbits and goats (ProteoGenix SAS, 67300 Schiltigheim, France).

IgY antibodies to the Hcrt pre-pro-peptide was generated by injecting a synthetic peptide, CVTTTALAPRGGSGV (Innovagen, Lund, Sweden), conjugated to KLH, into four hens. The N-terminal cysteine was added to allow for affinity-purification. IgY was purified from eggs and further affinity-purified using Pierce UltraLinkTM Iodoacetyl columns (Thermofisher) and eluted at pH7.0 using ActiSep (Sterogene). Peptide conjugation, IgY production and purification was conducted by Agrisera, Umea, Sweden.

### Confocal Imaging and Image Analysis

Fluorescent images were obtained with Leica DMi8 Widefield Inverted, Leica SP8 Point Scanning, Zeiss LSM900 Airyscan 2 or Zeiss AxioScan Z1 Fluorescent Imager microscopes. Confocal stacks were merged using Fiji software.^108^ Compilation of images and graphs was done in Adobe Illustrator. Microsoft Excel and GraphPad Prism were used for statistical analyses, data compilation and graphical representation. Quantification of immunostaining is available in Table S1.

### Single nuclei RNA-Seq

The embryonic or adult mouse hypothalamus was dissected out and nuclei were freshly prepared, while human hypothalamic organoids were frozen before nuclei preparation. Nuclei were isolated using a single nuclei isolation kit (cat no. PN1000494, 10xGenomics, Pleasanton, CA, USA). RNA-Seq libraries were generated using the Illumina Stranded mRNA Library Ligation kit (Illumina, 20040534) and Illumina RNA UD Indexes Ligation (Illumina, 20091657) according to the standard manufacturer’s protocol. Briefly, total RNA was enriched for mRNA using an oligo-dT bead isolation method, and the mRNA was fragmented in a heat fragmentation step. cDNA was synthesized from the fragmented RNA using random primers. The first strand cDNA was converted into dsDNA in the presence of dUTP to prevent subsequent amplification of the second strand and thus maintaining the strand orientation of the original mRNA. The 3’ ends of the cDNA were adenylated and pre-index anchors were ligated. The libraries were amplified with PCR, incorporating unique indexes for each sample to produce libraries ready for sequencing. The libraries were quantified on the Revvity LabChip GX Touch with the DNA High Sensitivity Reagent kit (Revvity, CLS760672). Libraries were pooled in equimolar ratios.

#### Sequencing data preprocessing

Sequencing was performed with 10× Genomics Chromium platform (Illumina NovaSeq X). After sequencing, fastq files were generated using bcl2fastq2 (v2.20.0.422), and sequencing adapters were trimmed. The demultiplexing, barcoded processing, gene counting and aggregation were made using the Cell Ranger software v10.0 (https://www.10xgenomics.com/support/software/cell-ranger/latest). Reads were mapped to mouse (mm10) and human (hg38) reference genomes to generate gene expression matrices. For nuclei numbers, UMI/nuclei and genes/nuclei detected, for all samples analysed, see Table S7.

#### Single-nuclei dataset processing and quality control

Raw 10x Genomics filtered feature–barcode matrices were imported and converted to Seurat objects. Genes detected in at least three cells and cells expressing at least 200 genes were retained. Mitochondrial transcript content was quantified using PercentageFeatureSet (pattern ^Mt-). Quality control was performed separately for each replicate by examining the distributions of nFeature_RNA, nCount_RNA, and percent.mt. Cells were retained when they met the following criteria: nFeature_RNA > 200, nCount_RNA > 200, percent.mt < 5%, and dataset-specific upper limits for nFeature_RNA (6,000-7,000 for mouse, 8,000 for HypoOrgs) and nCount_RNA (20,000-30,000 for mouse, 50,000 for HypoOrgs), which were applied to remove low-quality cells and likely multiplets. Each replicate was processed independently using Seurat’s standard workflow. Data were log-normalized and highly variable genes were identified and scaled. Dimensionality reduction was performed with PCA, followed by neighbourhood graph construction (dims = 1-50), clustering, and visualization using UMAP (dims = 1-50). Processing replicates separately allowed the quality and structure of each sample to be evaluated prior to integration.

#### Doublet detection

Doublets were identified separately for each replicate using DoubletFinder. Parameter optimization was performed and the optimal pK value was selected based on the maximum BCmetric. DoubletFinder was then run using PCs 1-50 with pN = 0.25 and dataset-specific pK values and expected doublet numbers. Only cells classified as singlets were retained for downstream analyses.

#### Replicate merging and Harmony integration

For each dataset, singlet-filtered replicates were merged, while genotype and replicate information were preserved in the metadata. The merged objects were reprocessed through normalization, PCA (dims = 1-50), neighbor graph construction, clustering, and UMAP embedding. To correct replicate-associated batch effects, Harmony integration was performed using SampleID as the batch variable. Downstream analyses were conducted in Harmony space, including neighbour graph construction and clustering (dims = 1-20 for mouse, 1-50 for HypoOrgs), followed by UMAP visualization using Harmony (dims = 1-20 for mouse, 1-50 for HypoOrgs).

#### Cell type annotation

For mouse, clusters were annotated manually based on canonical marker gene expression assessed using DotPlot and FeaturePlot. Major cell classes identified across datasets included GABAergic neurons, glutamatergic neurons, dopaminergic neurons, radial glia, neuronal precursors, oligodendrocyte lineage cells (OPCs, pre-myelinating and mature oligodendrocytes), astrocytes, endothelial cells, macrophages, fibroblasts, pericytes, and erythrocytes. For HypoOrgs, the top 50 marker genes per cluster were examined (FindAllMarkers) and interpreted in combination with established canonical markers of major brain cell types. Cell identities were assigned based on known gene expression signatures and previously published human hypothalamic and neuronal lineage datasets.

### Mouse sn-RNA-seq analysis

#### Pseudobulk generation

To account for biological replication in differential expression analyses, pseudobulk counts were generated from the Seurat objects. Raw RNA counts were extracted and summed per gene within each biological replicate. Sample identity and genotype (WT or cKO) were retained as metadata. Pseudobulk count matrices and associated metadata tables were exported for each dataset.

#### Differential expression analysis

Differential expression analysis was performed using the edgeR quasi-likelihood framework. DGEList objects were constructed from pseudobulk counts and normalized using calcNormFactors with TMM normalization. Genes were filtered using keepByExpr. Dispersion parameters were estimated and robust quasi-likelihood negative binomial models were fitted using glmQLFit. Differential testing between cKO and WT samples was conducted, with genotype modelled as a two-level factor referencing WT. All tested genes were reported, and statistical significance was defined as FDR ≤0.05. Genes with |log2FC| ≥0.25 and FDR ≤0.05 were considered differentially expressed.

#### Extraction of Lhx9/Rfx4 double-positive cells

For each annotated Seurat object, raw RNA counts were retrieved from all layers of the RNA assay and summed per gene for each cell. Gene positivity was defined as raw counts greater than zero. Cells were classified as Lhx9□, Rfx4□, or double-positive. Only cells co-expressing both genes (Lhx9>0 and Rfx4>0) were retained for subsequent analyses.

#### Integration of double-positive populations

Double-positive cells were analysed separately for embryonic and postnatal datasets. For embryonic analysis, E14_Tead1 and E14_Vgll2 cells were combined, whereas P20_Tead1 and P25_Vgll2 cells were merged for postnatal analysis. Merged datasets were normalized using SCTransform, followed by PCA. Batch effects were corrected using Harmony with the dataset label specified as the grouping variable. Harmony embeddings were computed using PCA dimensions 1–30. UMAP embeddings were generated from the Harmony reduction, and neighbour graphs were constructed in Harmony space. Clustering was performed using the Leiden algorithm (algorithm = 4) at resolution 0.6.

#### Marker visualization in double-positive populations

Expression of neuropeptide genes within the integrated double-positive populations was visualized using FeaturePlot. Hcrt expression was displayed on integrated UMAP embeddings and visualized across genotypes when relevant.

#### Hcrt-positive cell filtering and genotype comparison

For genotype-level comparisons of Hcrt expression, analyses were restricted to cells with Hcrt raw counts >0. In Seurat v5 objects containing layered RNA assays, layers were unified using JoinLayers when required to ensure consistent expression retrieval. Gene matching was performed in a case-insensitive manner. Genotype labels were standardized (WT, Tead1-cKO, Vgll2-cKO), and condition was treated as a factor with WT as the reference level. Expression distributions were visualized using violin plots generated from RNA assay expression values (layer = “data”), with jittered points representing individual cells.

### Human HCRT analysis

#### Cross-dataset integration

Single-cell datasets from adult hypothalamus, embryonic hypothalamus, and HypoOrgs were integrated using Seurat v5. Cells expressing HCRT were extracted from each dataset (HCRT>1 for adult and embryonic datasets, HCRT>0 for organoids). Metadata was standardized by assigning dataset labels (Adult, Embryonic, Organoid) and adding dataset-specific prefixes to ensure unique cell barcodes. Each data set was normalized using SCTransform. Integration features were selected using SelectIntegrationFeatures, and datasets were prepared using PrepSCTIntegration. PCA was performed for each dataset, and integration anchors were identified using FindIntegrationAnchors with SCT normalization and reciprocal PCA (reduction = “rpca”). Anchors were combined using IntegrateData. The integrated object was analysed using PCA and UMAP (dims = 1-50), followed by neighbour graph construction (FindNeighbors) and clustering (FindClusters, resolution = 0.8). UMAP visualizations were generated using DimPlot.

#### Gene expression visualization

Marker gene expression (including HCRT, VGLL2, and TEAD1) was visualized using FeaturePlot. Because RNA assays contained multiple count layers, per-cell expression vectors were constructed by extracting gene counts from each layer and storing log1p-transformed values in metadata. Dataset-specific feature plots were also generated by subsetting the integrated object according to dataset labels.

#### Cluster marker identification

Cluster marker genes were identified using Seurat’s FindMarkers. Markers were detected using the Wilcoxon rank-sum test (test.use = “wilcox”), considering only positive markers (only.pos = TRUE) with thresholds of min.pct = 0.25 and logfc.threshold = 0.25. Marker tables and the top 100 markers per cluster were exported.

#### Gene Ontology analysis

Gene Ontology enrichment analyses were summarized across clusters for BP, CC, and MF categories. Significant terms were defined as those with adjusted p-value ≤0.05 and gene count >5. Within each cluster, the top 10 terms ranked by GeneRatio were selected. Enrichment results were visualized using dot plots generated with ggplot2, where color represented −log10(p.adjust).

#### HypoOrgs EN21 analysis

LHX9/RFX4 double-positive cells were identified by extracting raw RNA counts and defining gene positivity as counts greater than zero. Cells co-expressing both genes were subsetted for downstream analysis. The subsetted dataset was normalized using SCTransform, followed by PCA. Batch effects were corrected using Harmony with the appropriate metadata variable as the grouping factor. Harmony embeddings were calculated using PCA dimensions 1–30. UMAP embeddings were generated from the Harmony reduction, followed by neighbour graph construction and clustering using the Leiden algorithm (algorithm = 4, resolution = 0.6). Clusters were stored as seurat_clusters, and transcriptional structure was visualized using UMAP.

### Maintenance of human iPSC

A human iPSC line with a doxycycline (dox)-inducible dCas9-VPR cassette, including a constitutive TRE activator rtTA transgene, targeted to the AAVS1 safe harbor site,^89^ was obtained from Ernst Wolvetang (The University of Queensland, Australia). hiPSCs were maintained at 37°C with 5% CO in tissue culture incubators and cultured on Matrigel-coated 6-well plates (Falcon Matrigel hESC-qualified Matrix, LDEV-free; Corning, Cat. #354277). Each well contained 2 mL of mTeSR Plus Medium (Stemcell Technologies, Cat. #1000276), with complete medium changes performed every other day. When cultures reached 70-80% confluence, cells were passaged at a 1:20 split ratio using ReLeSR (Stemcell Technologies, Cat. #1000483). Briefly, medium was aspirated, cells were washed with 1 mL DPBS (Gibco, Cat. #14190144) and incubated with a thin layer of ReLeSR for 5 min at 37°C, 5% CO□, with occasional manual shaking. Cells were then gently pipetted 3-5 times to dissociate into small clumps and replated at a 1:20 ratio into new Matrigel-coated 6-well plates with mTeSR Plus medium. Experiments involving hiPSCs were performed with the approval of the University of Queensland Human Research Ethics Committee.

### Generation of HypoOrgs

hiPSCs (60-70% confluent) were dissociated into single cell suspension by incubating with Accutase (Gibco, Cat. #A1110501) for 3-5 mins at 37°C, resuspended in mTeSR Plus medium supplemented with 10µM Y-27632 (Sigma-Aldrich #Y0503-1MG) and seeded at a density of 1×10 cells per well into no-coating ultra-low attachment 96-well U-bottom plates (Nunclon Sphera, Thermo Scientific #174925). Plates were centrifuged at 200g for 3 min to combine cells together into a spheroid and then transferred to an incubator (37°C, 5%CO_2_) for subsequent culture. The following day (Day 0; D0), spheroids were washed twice with DPBS to remove Y-27632 and cultured in KnockOut Serum Replacement (KSR)* medium; DMEM/F12 (Gibco #11320033), 20% KnockOut™ Serum Replacement (Gibco #10828028), 2mM/1× GlutaMAX (Gibco #350500061), 0.1mM/1× MEM NEAA (Minimum Essential Medium Non-Essential Amino Acids Solution, MEM NEAA, Stemcell Technologies #07600), 100U/ml PenStrep (Gibco #15140122), 0.1mM 2-Mercaptoethanol (Gibco #21985023), supplemented with 200nM retinoic acid.^109^ After 24 hours (D1), spheroids were washed twice with DPBS to remove retinoic acid and cultured in fresh KSR* medium. On D2, the full medium was changed to fresh KSR* medium, with the addition of 100nM LDN193189 (Stemcell Technologies #72147), 10μM SB431542 (Stemcell Technologies #72234), and 2μM XAV939 (Stemcell Technologies #72674). From D4, KSR* medium was gradually replaced with N2* medium; Neurobasal™ Medium (Gibco #21103049) supplemented with 1×N2 Supplement (Thermo Scientific #17502048), 2mM/1× GlutaMAX, 0.1mM/1× MEM NEAA, 100U/ml PenStrep, every other day until D10 to 100% N2* medium, and simultaneously, the concentrations of LDN193189, SB431542, and XAV939 were also gradually decreased. 1μM SAG (Smoothened agonist, SAG; Stemcell Technologies #73412), and 1μM Purmorphamine (Stemcell Technologies #1001049) were added from D2 to D8.^70^ 2μM AKT Inhibitor VIII (Stemcell Technologies #72942) was added from D4 to D14.^110^ 5μM DAPT (Stemcell Technologies #72082) was added from D8 to D14.^111^ From D16 onwards, cells were fully switched into maturation medium*; Neurobasal™-A Medium (Gibco #10888022), 2mM/1× GlutaMAX™ Supplement (Gibco #350500061), 1× N2 Supplement (Thermo Scientific #17502048), 1× B27 Supplement (Thermo Scientific #17502044), 0.075% Sodium Bicarbonate (Gibco #25080094), 200nM Ascorbic Acid (Rowe Scientific PTY LTD #CL0190), 1μM dibutyryl cyclic AMP (dcAMP) (Sigma-Aldrich #D00627-100MG), 10ng/ml GDNF (Stemcell Technologies #78139), 10ng/ml BDNF (Stemcell Technologies #78133), 10ng/ml CNTF (Stemcell Technologies #78170) 2mM/1× GlutaMAX, 0.1mM/1× MEM NEAA, 100U/ml PenStrep. Cells were maintained in maturation medium by half-volume medium changes every other day. 2µg/ml doxycycline was continuously added at either D4 or D10 onward, to assess the impact of VGLL2 and TEAD1 activation on hypothalamic fate specification and the generation of HCRT and NPVF neurons.

### Lentivirus Infection

Lentiviral DNA constructs (Suppl Info 1) and lentiviral production were provided by VectorBuilder Inc (Chicago, IL, USA). hiPSCs were maintained in mTeSR Plus medium in 6-well plates. When cultures reached 30-40% confluency, lentiviral vectors were added to the medium at a multiplicity of infection (MOI) of 10, together with polybrene (10µg/ml) to enhance infection efficiency. Cells were exposed to the lentiviral vectors for 48 hours, after which the virus-containing medium was removed and replaced with fresh medium.

### FACS sorting

Infected hiPSCs were dissociated into single cells by incubating with Accutase for 3-5 mins at 37°C. The reaction was neutralized with mTeSR Plus medium, and cells were gently pipetted 3-5 times to obtain a single-cell suspension. The suspension was centrifuged at 300g for 5 mins, the supernatant was removed, and the pellet was resuspended in FACS buffer, consisting of 1xPBS, 1% Penicillin-Streptomycin, 0.1% BSA solution and 10mM Y-27632. The suspension filtered through a 40μm strainer into a 5ml Falcon round-bottom tube. Cell sorting was performed on a BD FACSAria Cell Sorter (BD Biosciences) to enrich for GFP and/or mCherry cells for subsequent culture. The gating strategy included: (1) P1 gate to exclude cell debris, (2) singlet gating to remove doublets, and (3) viability gating to exclude dead cells. The GFP and mCherry gates were defined based on control cells.

## Figure Legends

**Figure S1. VGLL and TEAD expression in the developing mouse hypothalamus.**

**(A-F)** Gene expression of *Tead2*, *Tead3, Tead4*, *Vgll1*, *Vgll3* and *Vgll4* in the developing mouse hypothalamus at E13.5, based upon sagittal ABA *in situ* hybridization^78^ or transverse spatial RNA-seq data.^30^ Green brackets and boxes depict the putative pT1-1/EN21 domain. Blue and red dashed lines depict the estimated level of sections in the spatial RNA-seq data when compared to the sagittal *in situ* hybridization images.

**Figure S2. Patterning gene expression in the mouse tuberal hypothalamus.**

**(A-O)** Gene expression of patterning and postmitotic TF genes in the developing mouse hypothalamus, based upon sagittal ABA *in situ* hybridization^78^ or transverse spatial RNA-seq data.^30^ Green brackets and boxes depict the putative pT1-1/EN21 domain. Blue and red dashed lines depict the estimated level of sections in the transverse spatial RNA-seq data when compared to the sagittal *in situ* hybridization images.

**Figure S3. TF expression in the mouse EN21/Hcrt/Npvf cluster.**

**(A)** UMAP clustering of sn-RNA-seq data from E10 mouse hypothalamus,^30^ showing expression of *Tead1* and *Rax* in many progenitor cells. AS, astrocytes; BC, blood cells; EC, ependymal cells; EnC, endothelial cells; GABA, GABAergic neurons; GLU, glutamatergic neurons; IPC1/2, intermediate progenitor cells; MG, microglia; OD, oligodendrocytes; OPC, oligodendrocyte precursor cells; RGC, radial glial cells; TC, tanycytes; VLMC, vascular and leptomeningeal cells. **(B)** UMAP clustering of sn-RNA-seq data from E14 mouse hypothalamus neurons,^30^ showing expression of *Hcrt*, *Npvf*, *Vgll2*, *Tead1*, *Rfx4*, *Lhx9* and *Hmx2* in the EN21 cluster. **(C-E)** Dot plots depicting expression of TF genes, *Hcrt* and *Npvf* in progenitors (C), the EN21 neuron cluster (D-E) during development^30^ and the adult C66-18 cluster.^5^

**Figure S4. *Vgll2* expression in the mouse EN7 and EN8 glutamatergic clusters.**

**(A-C)** UMAPs based upon sn-RNA-seq data from E14 mouse hypothalamus neurons,^30^ showing the EN7 and EN8 glutamatergic clusters. **(D-E)** UMAPs showing expression of *Vgll2* in the entire cell set (D) and selectively in EN7 (E) and EN8 (F) clusters.

**Figure S5. TF expression in the human EN21/HCRT/NPVF cluster.**

**(A)** UMAP clustering of sn-RNA-seq data from PCW5 human hypothalamus,^30^ showing expression of TEAD1 and RAX in many progenitor cells. AS, astrocytes; BC, blood cells; EC, ependymal cells; EnC, endothelial cells; GABA, GABAergic neurons; GLU, glutamatergic neurons; IPC1/2, intermediate progenitor cells; MG, microglia; OD, oligodendrocytes; OPC, oligodendrocyte precursor cells; RGC, radial glial cells; TC, tanycytes; VLMC, vascular and leptomeningeal cells. **(B)** UMAP clustering of sn-RNA-seq data from PCW13 human hypothalamus neurons^30^, showing expression of *HCRT*, *NPVF*, *VGLL2*, *TEAD1*, *RFX4*, *LHX9* and *HMX2* in the EN21 cluster. **(C-E)** Dot plots depicting expression of TF genes, *HCRT* and *NPVF* in progenitors (C), the EN21 neuron cluster (D-E) during development.^30^

**Figure S6. *Vgll2^cKO^*and *Tead1^cKO^* mouse mutants show loss of Hcrt and Npvf neurons at P0. (A-H)** Hcrt and Npvf expression in the P0 mouse hypothalamus in control, *Vgll2^cKO^*and *Tead1^cKO^*. (**I-L**) Quantification of Hcrt and Npvf cells in the entire P0 mouse brain by serial sectioning, showing significant reduction of Hcrt and Npvf cells (Student’s two-tailed T-test; mean+/-SD; ** = p<0.01, *** =p≤0.001, **** = p≤0.0001; n = 3 brains analysed). Scale bars = 200μm.

**Figure S7. *Vgll2* and *Tead1* genetically interact in the mouse hypothalamus.**

**(A-D)** Hcrt and Npvf expression in the P20 mouse hypothalamus in control (*Vgll2^fl/+^*; *Tead1^fl/+^*) and *Vgll2*; *Tead1* trans-heterozygotes (*Vgll2^fl/+^*; *Tead1^fl^; Sox1-Cre/+*). (**E-F**) Quantification of Hcrt and Npvf cells in the entire P20 mouse brain by serial sectioning, showing significant reduction of Hcrt and Npvf cells (Student’s two-tailed T-test; mean+/-SD; * = p<0.05, ** = p<0.01, *** =p≤0.001, **** = p≤0.0001; n = 3 brains analysed). Scale bars = 200μm.

**Figure S8. Vgll2 and Tead1 mouse mutants do not affect major hypothalamus cell types.**

**(A)** Dot plot for the top marker genes for the different major cell clusters in control and mutants, based upon mouse sn-RNA-seq data. Control and mutant were co-analysed at each stage. **(B)** Cell proportions of major cell types in control and mutants at different stages.

**Figure S9. Cell annotation and cell proportions for human HypoOrgs. (A)** Dot plot for the top marker genes for the different major cell clusters in control HypoOrgs. **(B)** Cell proportions of major cell types in control and TF misexpression.

**Figure S10. Generation of VGLL2 and TEAD1 expressing hiPSC lines. (A-F’’)** Control and infected hiPSCs 48 hours after infection with lentiviruses expressing VGLL2 (green) or TEAD1 (green), followed by 24 hours Dox induction and stained for DAPI (blue). Negative controls, without virus or without Dox, show no specific immunostaining, while positive controls show nuclear/cytoplasmic staining for VGLL2 and nuclear staining for TEAD1. **(G-I)** hiPSC lines after double rounds of FACS sorting for VGLL2, TEAD1 and VGLL2-TEAD1 lentivirus insertions, 24 hours after Dox induction, immunostained for VGLL2 (green) and TEAD1 (red) showing that most cells express these TFs. **(G-M)** Expression of VGLL2 alone resulted in both cytoplasmic and nuclear staining, whereas VGLL2 co-expression with TEAD1 resulted in an apparent increase in VGLL2 nuclear staining.

**Figure S11. VGLL2-TEAD1 co-misexpression in human hypothalamic organoids induces HCRT neuron differentiation.**

**(A-P)** Staining for HCRT and DAPI in 20μm sections of HypoOrgs at D40, with dox induction at D4 **(A-H)** and at D80, with dox induction at D10 **(I-P)**.

**Figure S12. GO analysis of human HCRT neuron clusters; Biological Processes.**

GO term enrichment for Top10 terms for human HCRT clusters, Biological Processes. Terms relevant to nervous system function are highlighted in red.

**Figure S13. GO analysis of human HCRT neuron clusters; Cellular Component.**

GO term enrichment for Top10 terms for human HCRT clusters, Cellular Component. Terms relevant to nervous system function are highlighted in red.

**Figure S14. GO analysis of human HCRT neuron clusters; Molecular Function.**

GO term enrichment for Top10 terms for human HCRT clusters, Molecular Function. Terms relevant to nervous system function are highlighted in red.

**Table S1. Quantification of mouse immunostaining.**

**Table S2. Top-50 marker genes for mouse cell clustering.**

**Table S3. DEGs in mouse *Vgll2^cKO^* and *Tead1^cKO^* mutants based upon sn-RNA-seq.**

**Table S4. Top-50 marker genes for HypoOrgs cell clustering.**

**Table S5. DEGs in human HypoOrgs based upon sn-RNA-seq.**

**Table S6. Human HCRT sub-cluster marker genes and GO enrichment. Table S7. sn-RNA-seq nuclei numbers and statistics.**

**Supplemental Info 1. DNA sequences.**

(A) DNA sequences used to generate mouse Npvf and Vgll2 GST fusion proteins in *E. coli* for antibody production. (B). DNA sequences and lentiviral vector maps for human TEAD1 and VGLL2 expression in human hypothalamic organoids.

