## Supplemental Figures for "Vgll2 and Tead1 Govern Generation of Mouse and Human Hypothalamic Hypocretin (Orexin) Neurons"

### Supplemental Figure 1

#### VGLL and TEAD expression in the mouse hypothalamus

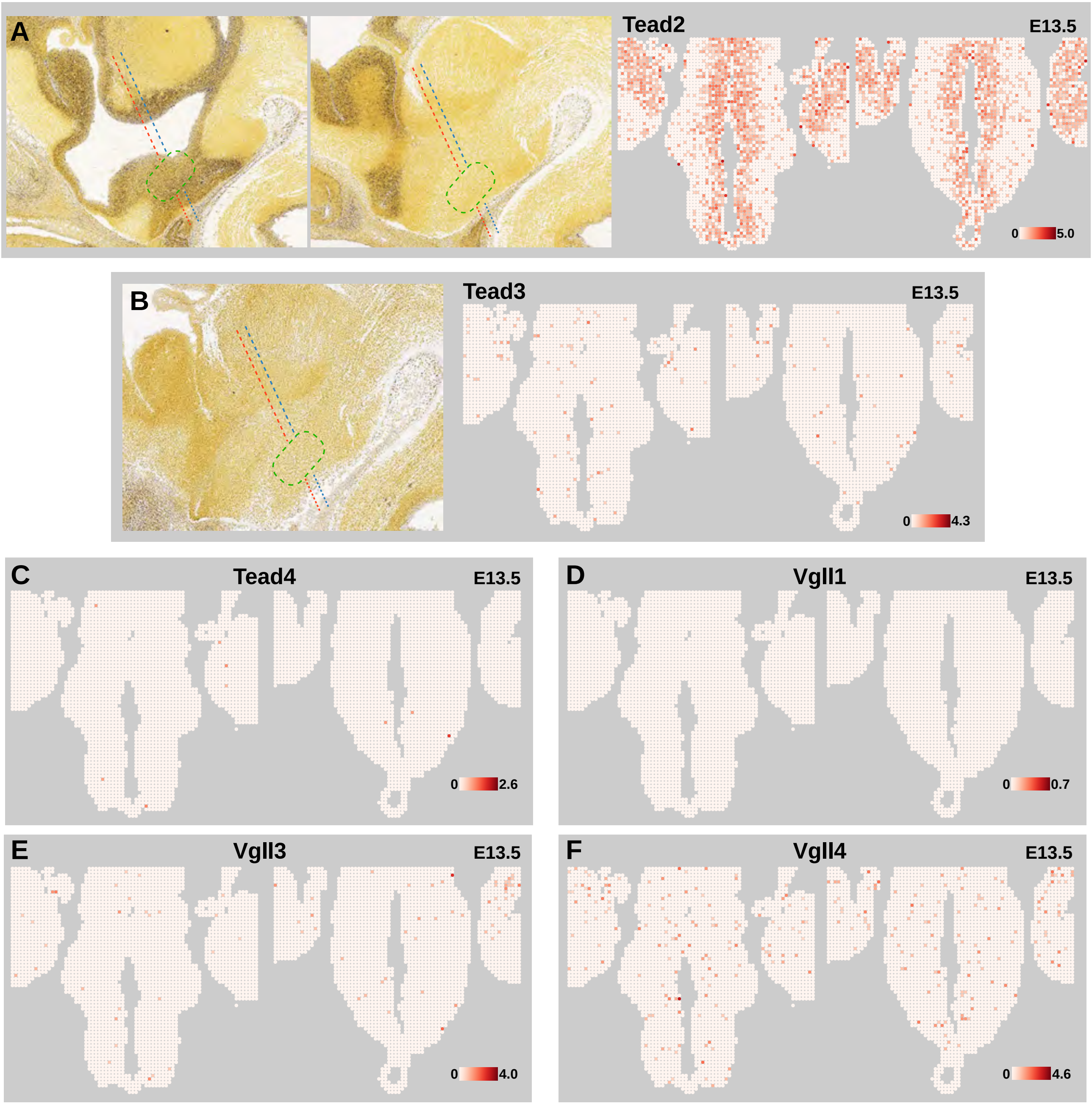

### Supplemental Figure 2

#### Patterning of the tuberal hypothalamus

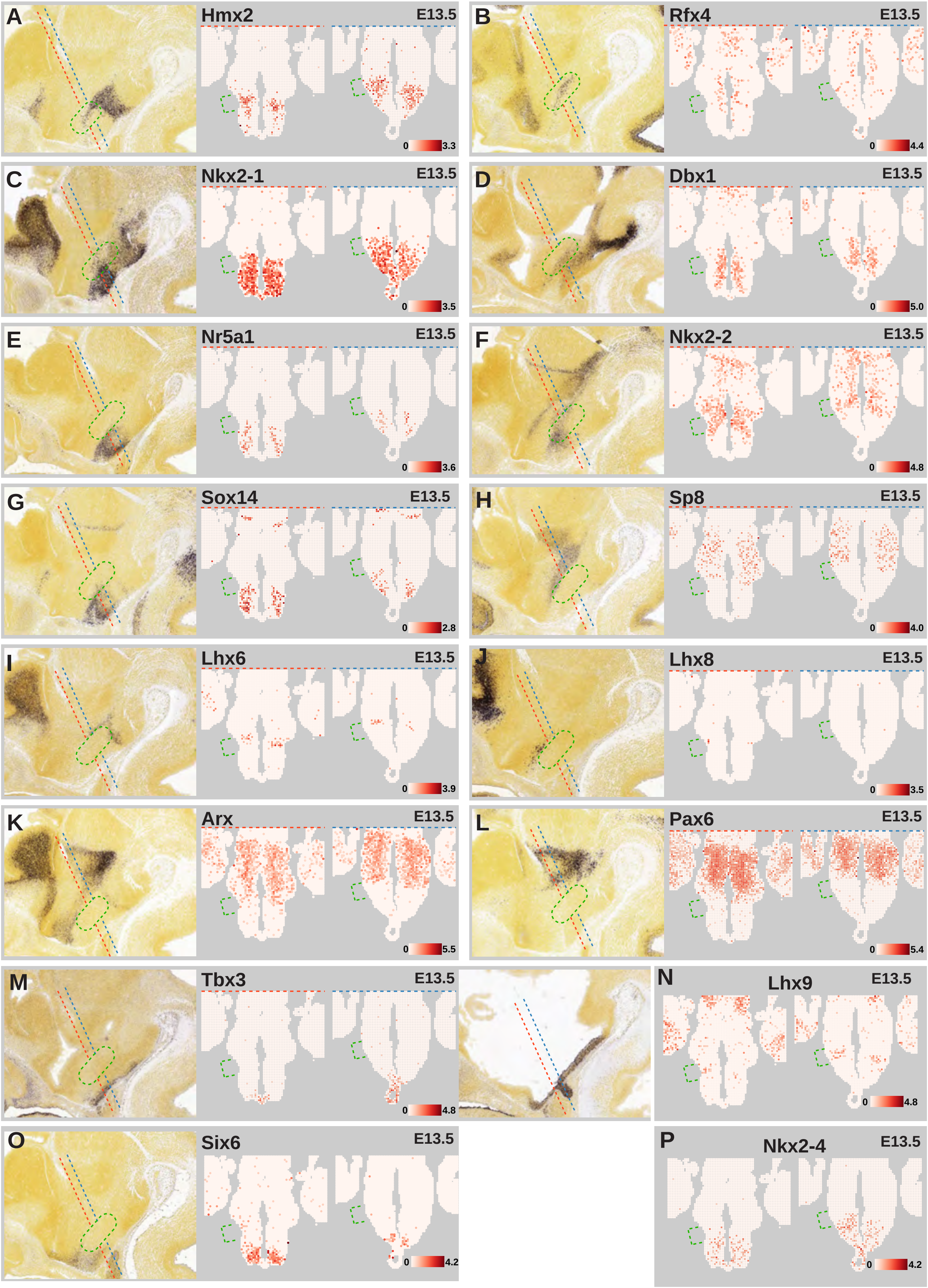

### Supplemental Figure 3

#### TF expression in the mouse EN21/Hcrt/Npvf cluster

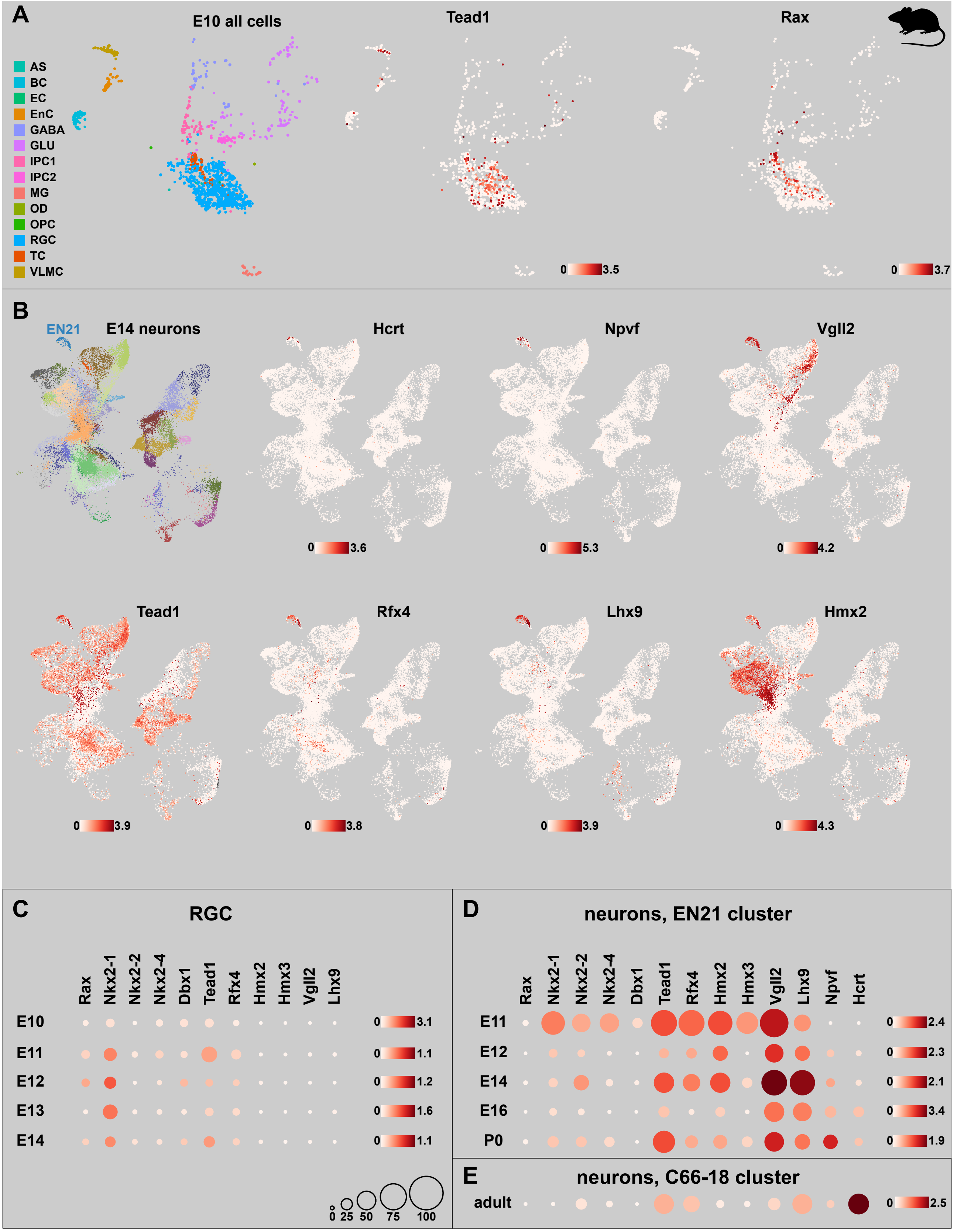

### Supplemental Figure 4

#### *Vgll2* expression in mouse EN7 and EN8 clusters

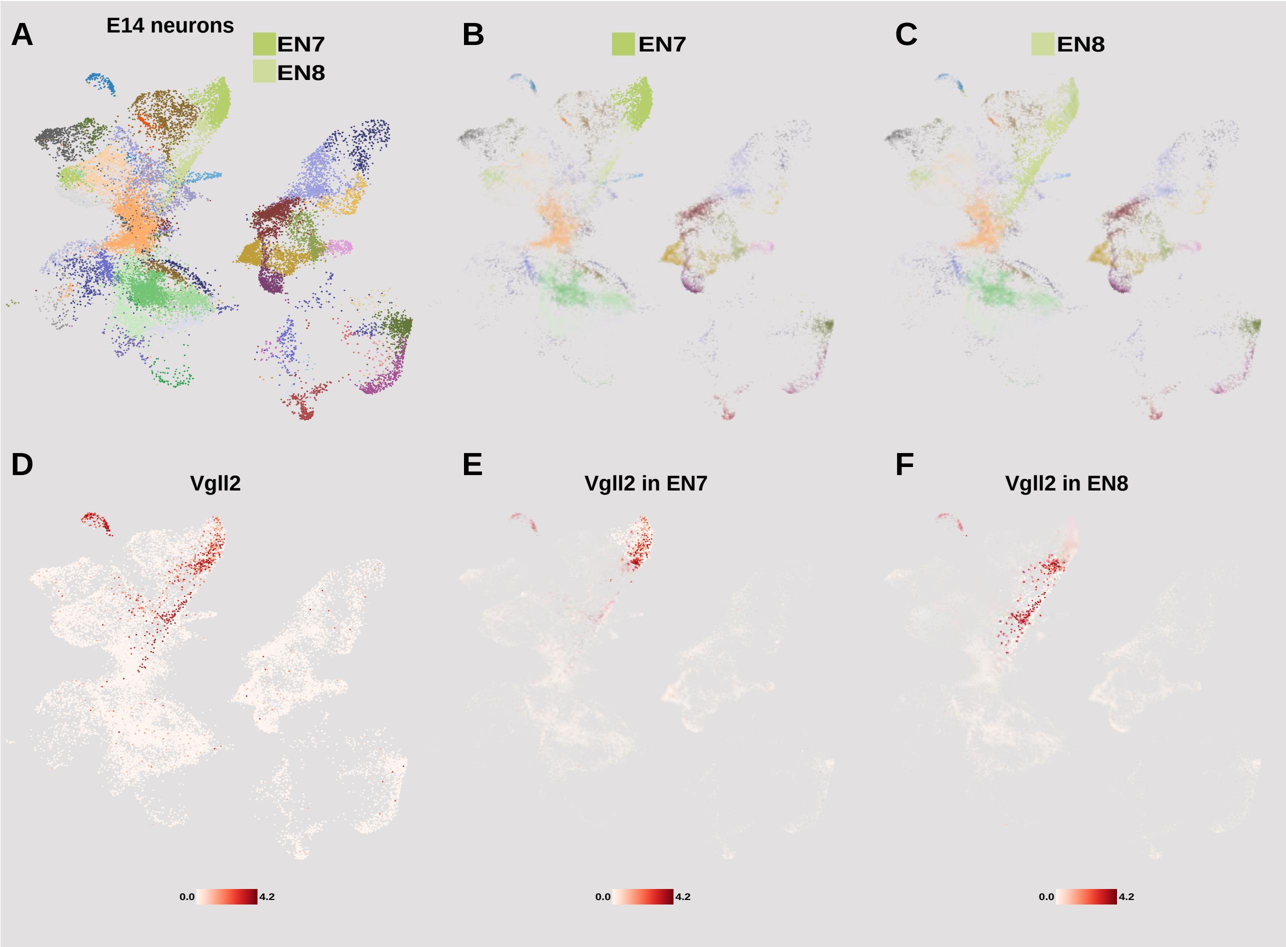

### Supplemental Figure 5

#### TF expression in the human EN21/HCRT/NPVF cluster

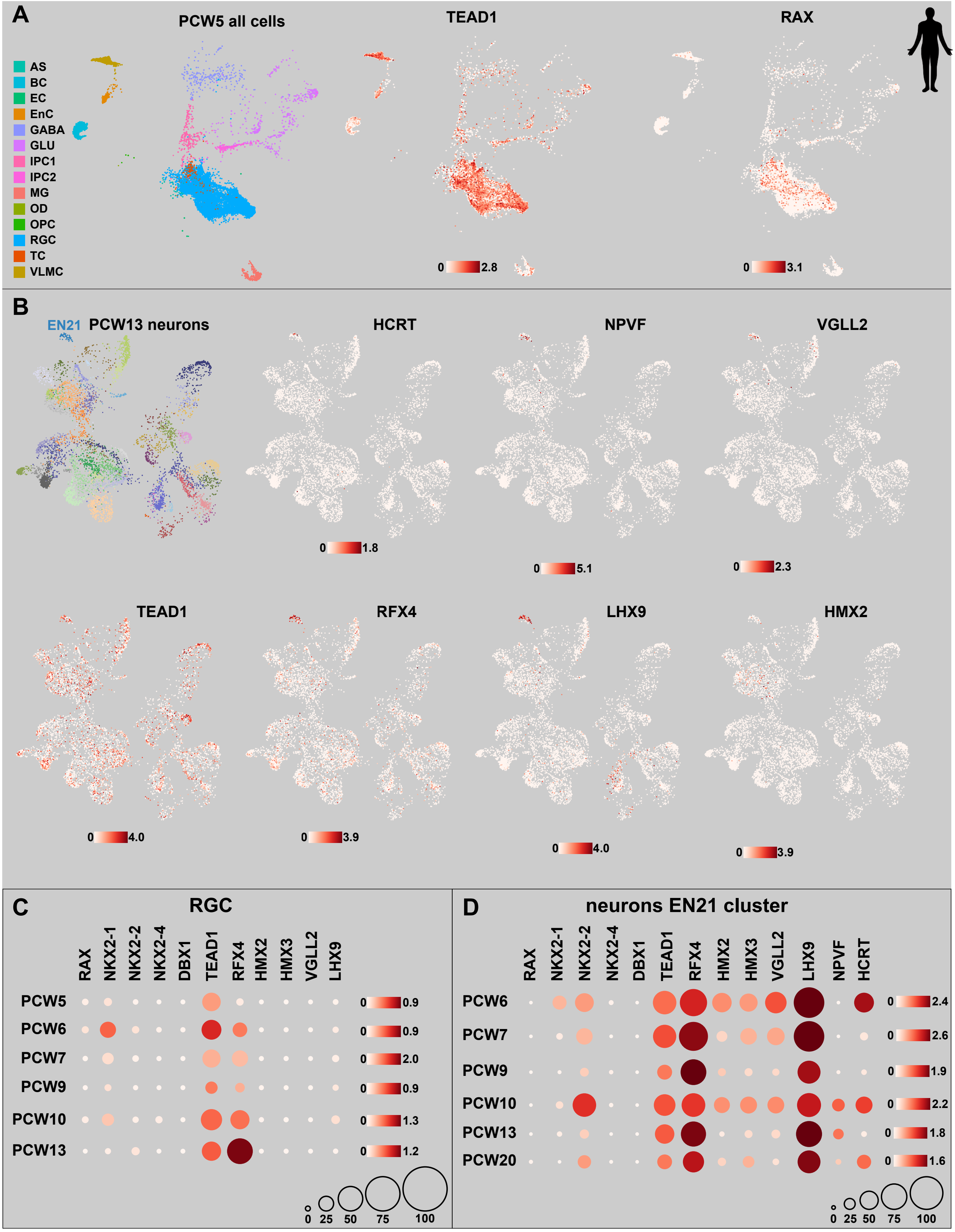

### Supplemental Figure 6

#### *Vgll2* and *Tead1* mutants affect Hcrt and Npvf at P0

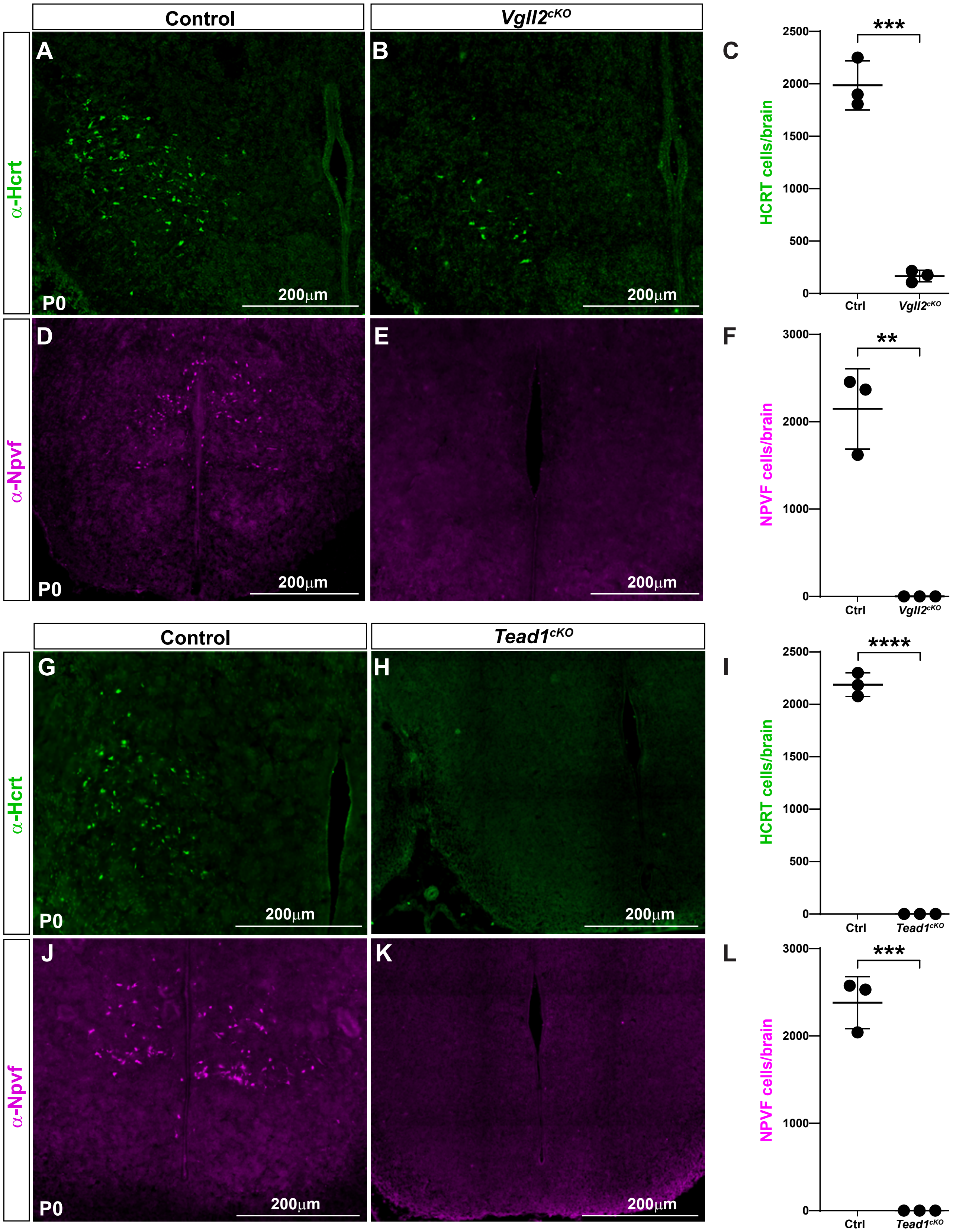

### Supplemental Figure 7

#### *Vgll2* and *Tead1* genetically interact

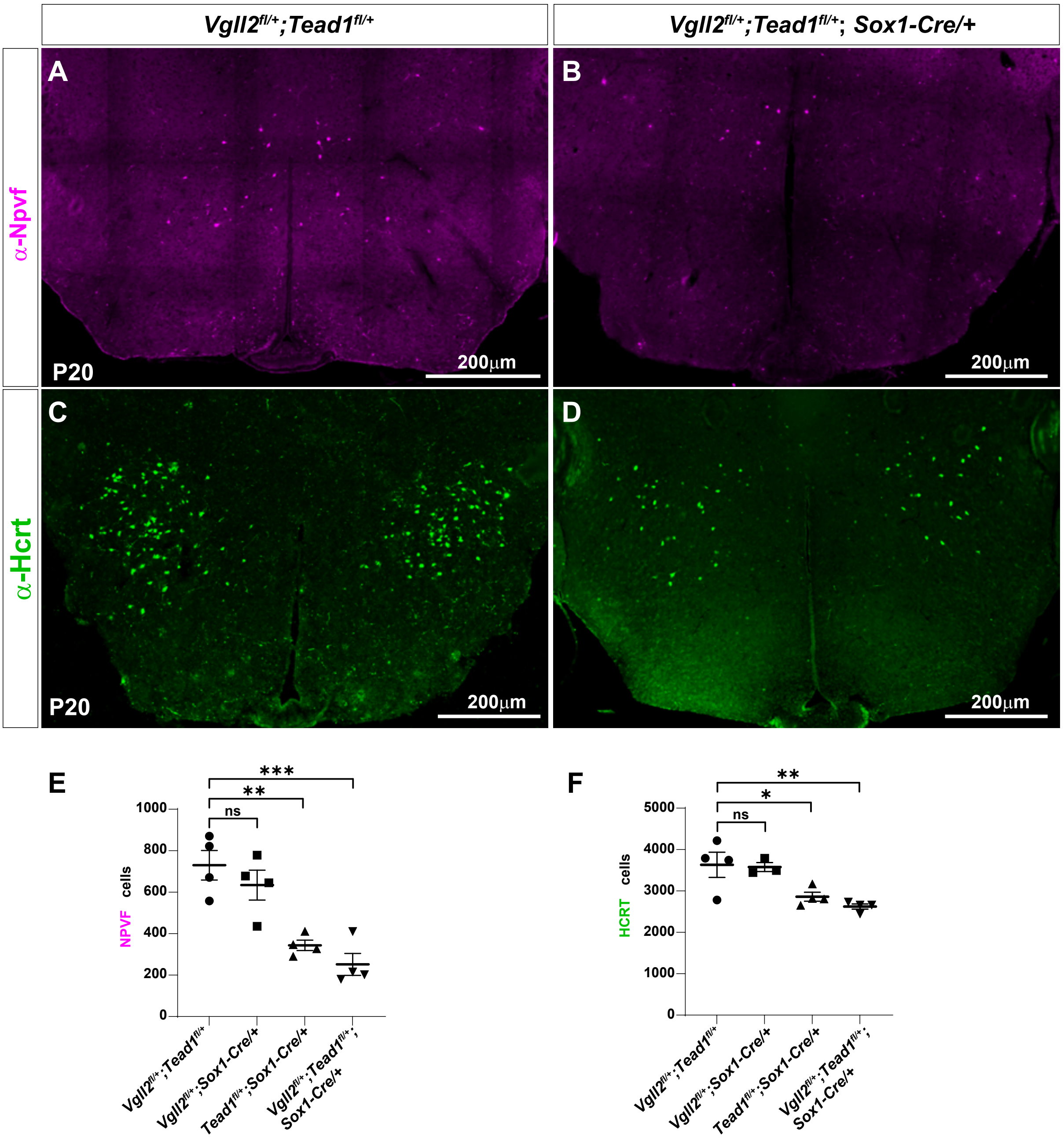

### Supplemental Figure 8

#### *Vgll2* and *Tead1 mouse* mutants do not affect major cell types

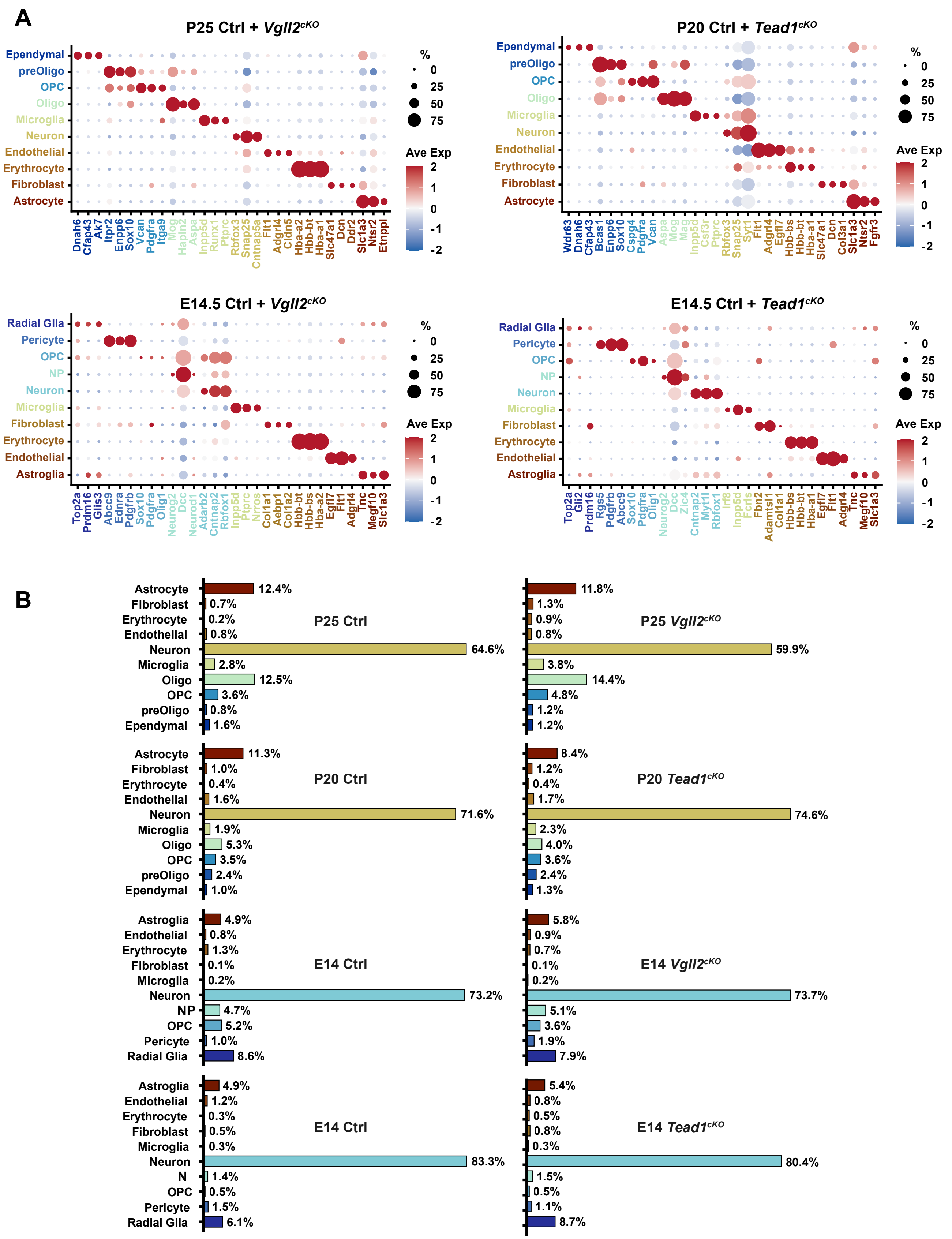

### Supplemental Figure 9

#### Cell annotation and cell proportions of human HypoOrgs

A

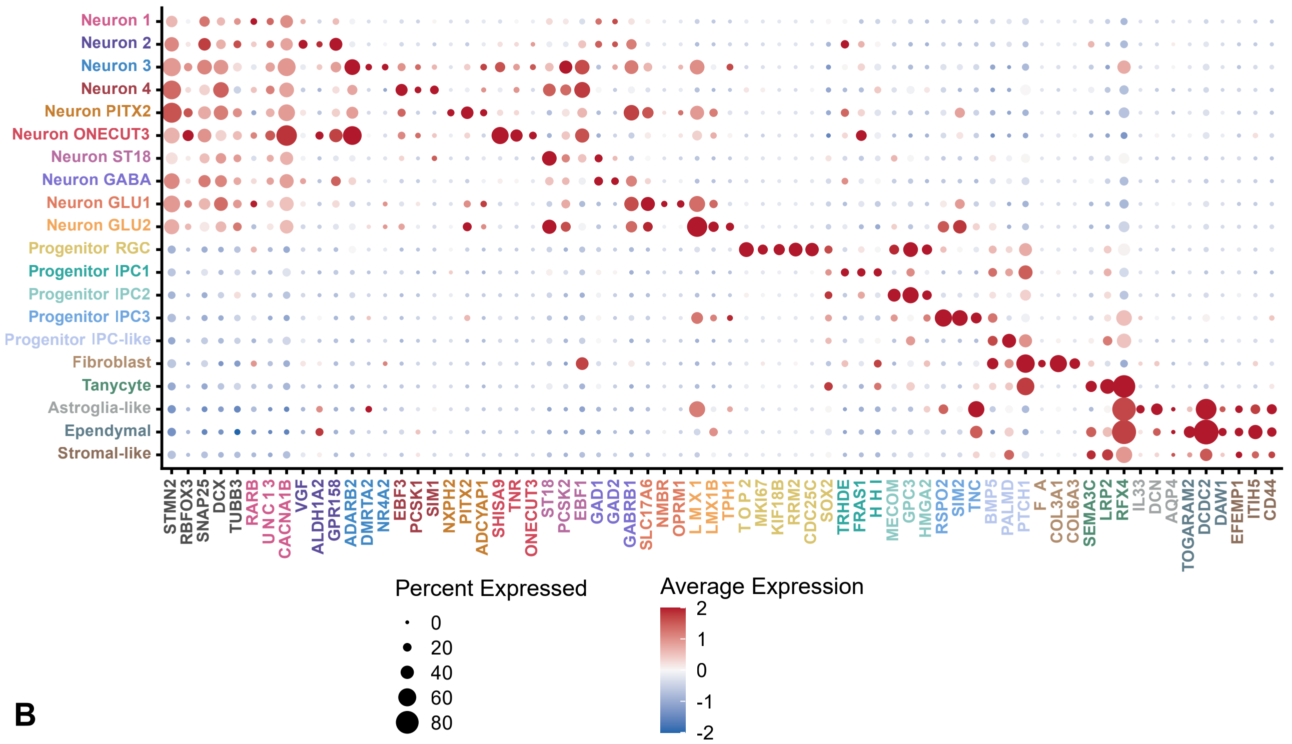


B

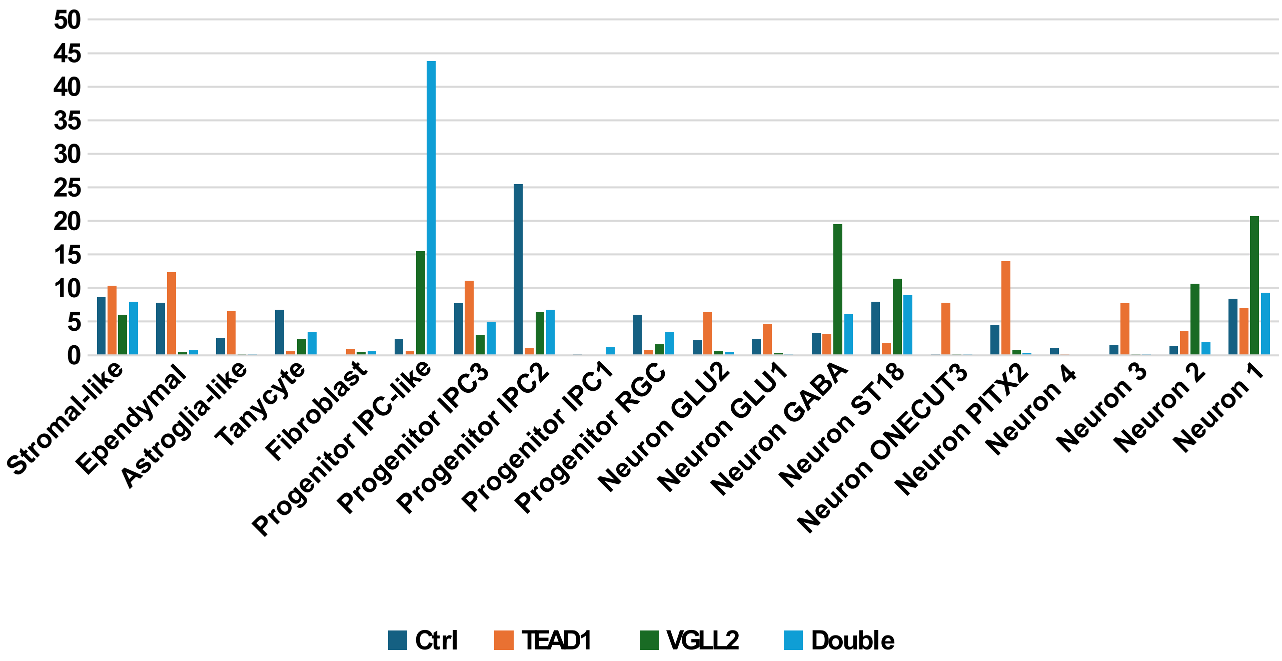

### Supplemental Figure 10

#### Generation of *VGLL2* and *TEAD1* expressing hiPSC lines

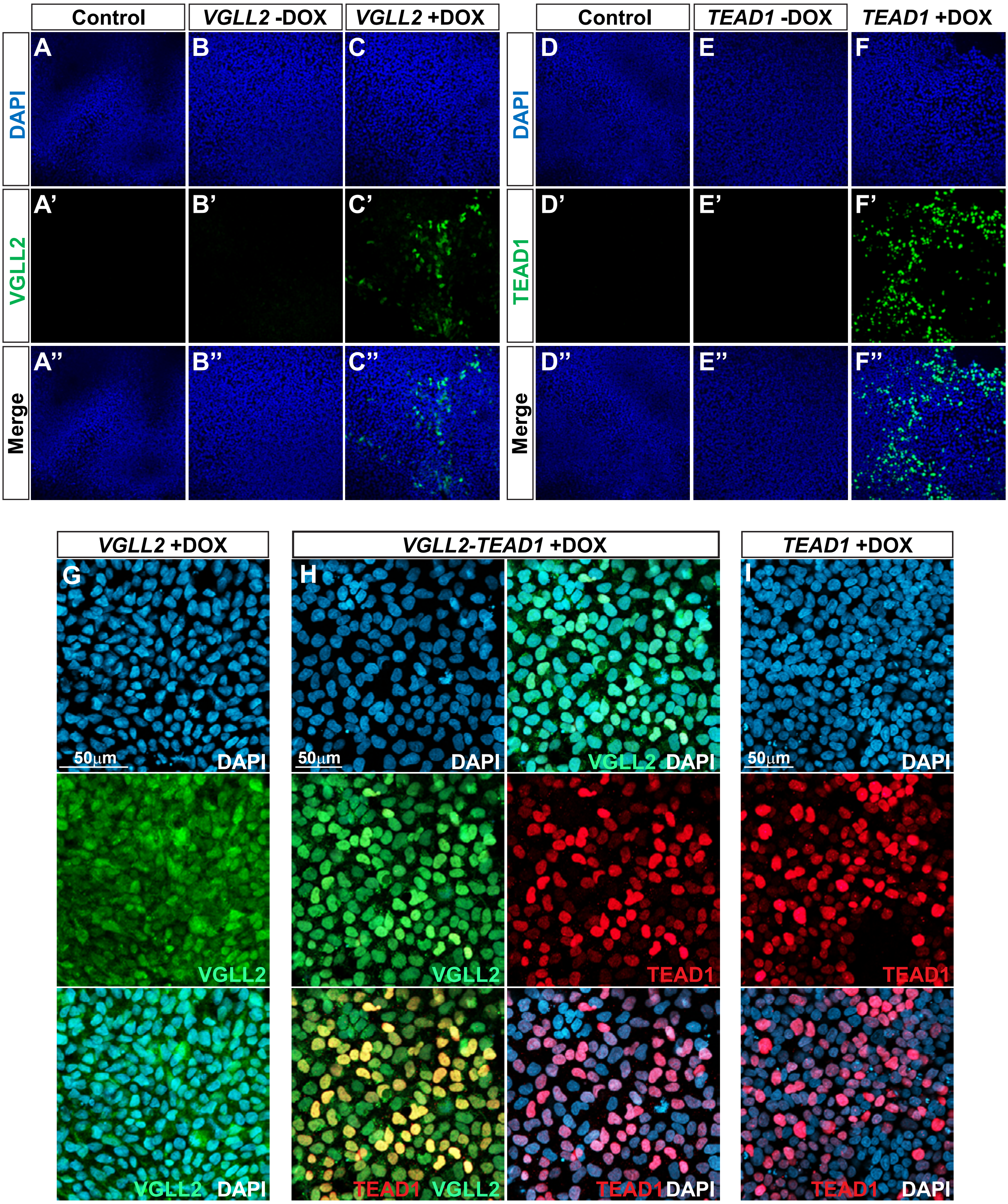

### Supplemental Figure 11

#### Co-misexpression of *VGLL2-TEAD1* triggers extra HCRT cells

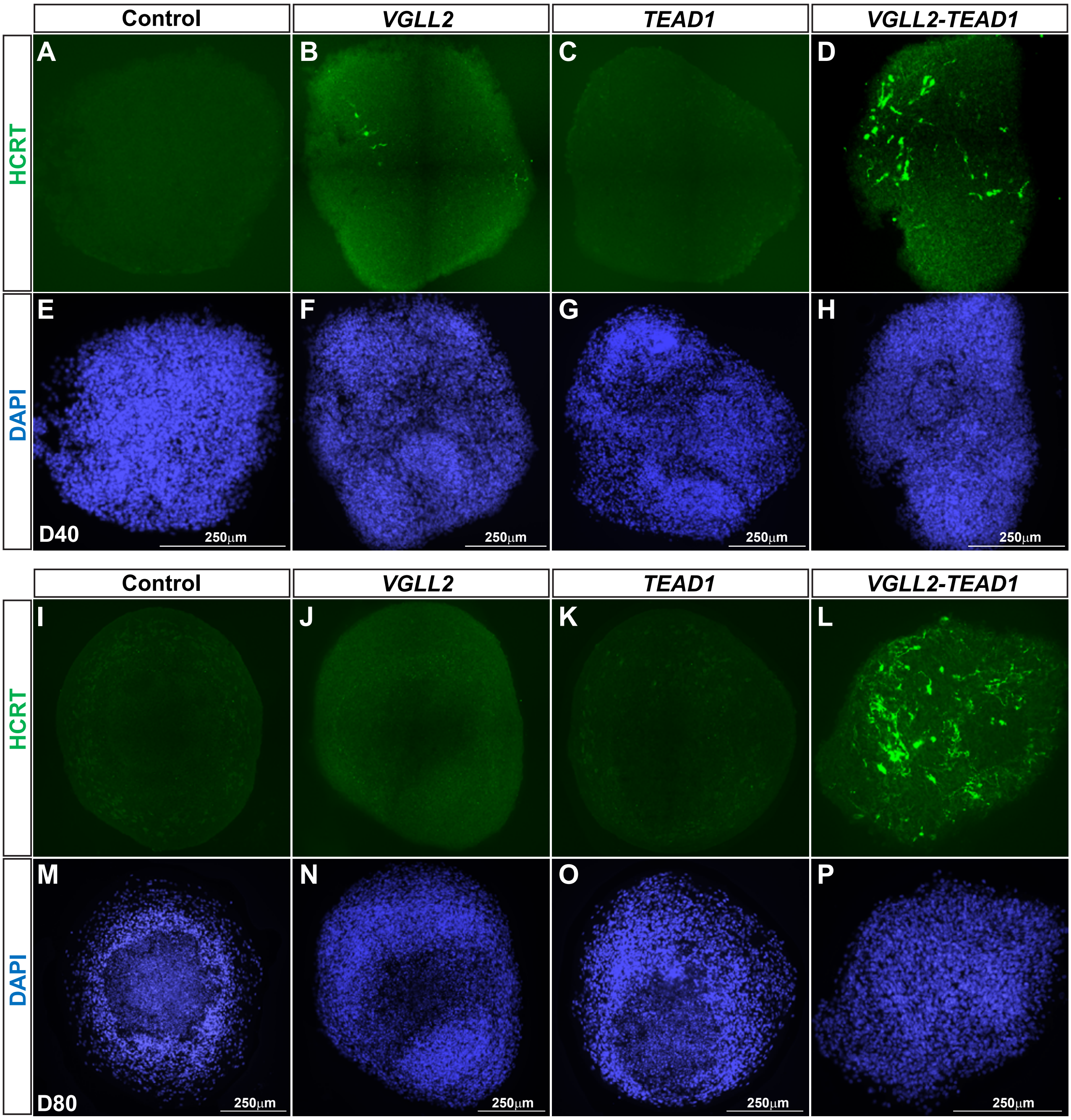

### Supplemental Figure 12

#### GO analysis (BP) of human HCRT cluster markers

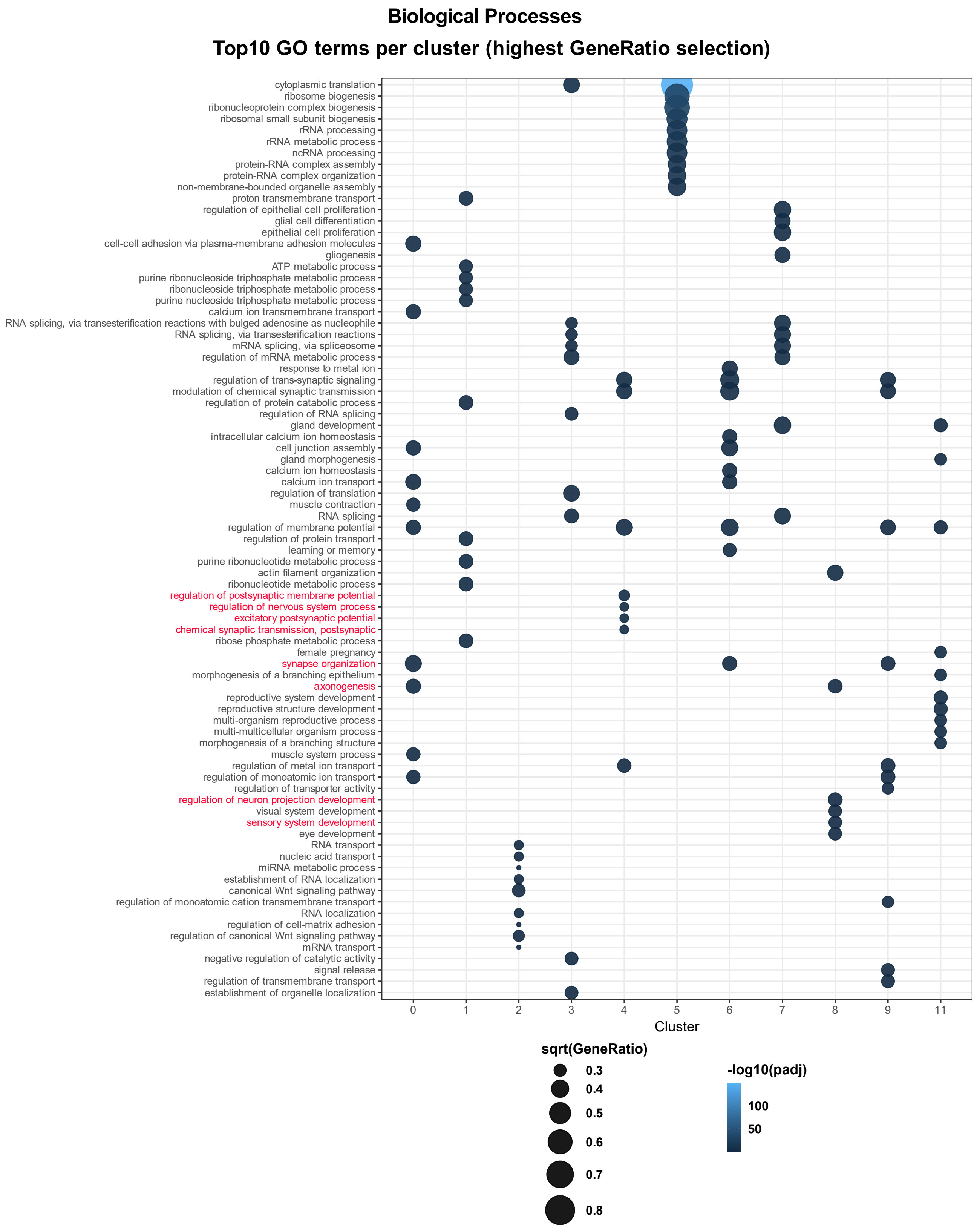

### Supplemental Figure 13

#### GO analysis (CC) of human HCRT cluster markers

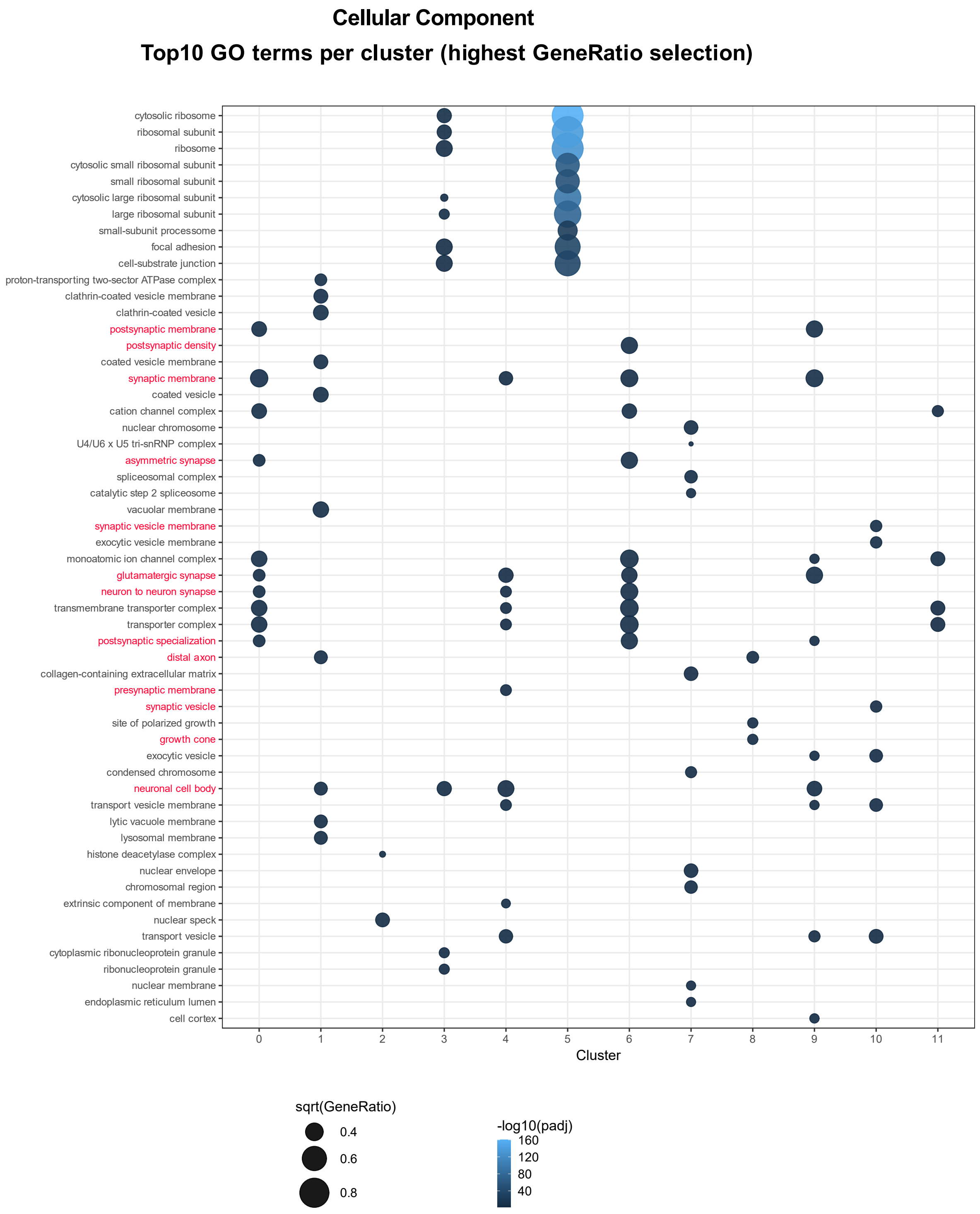

### Supplemental Figure 14

#### GO analysis (MF) of human HCRT cluster markers

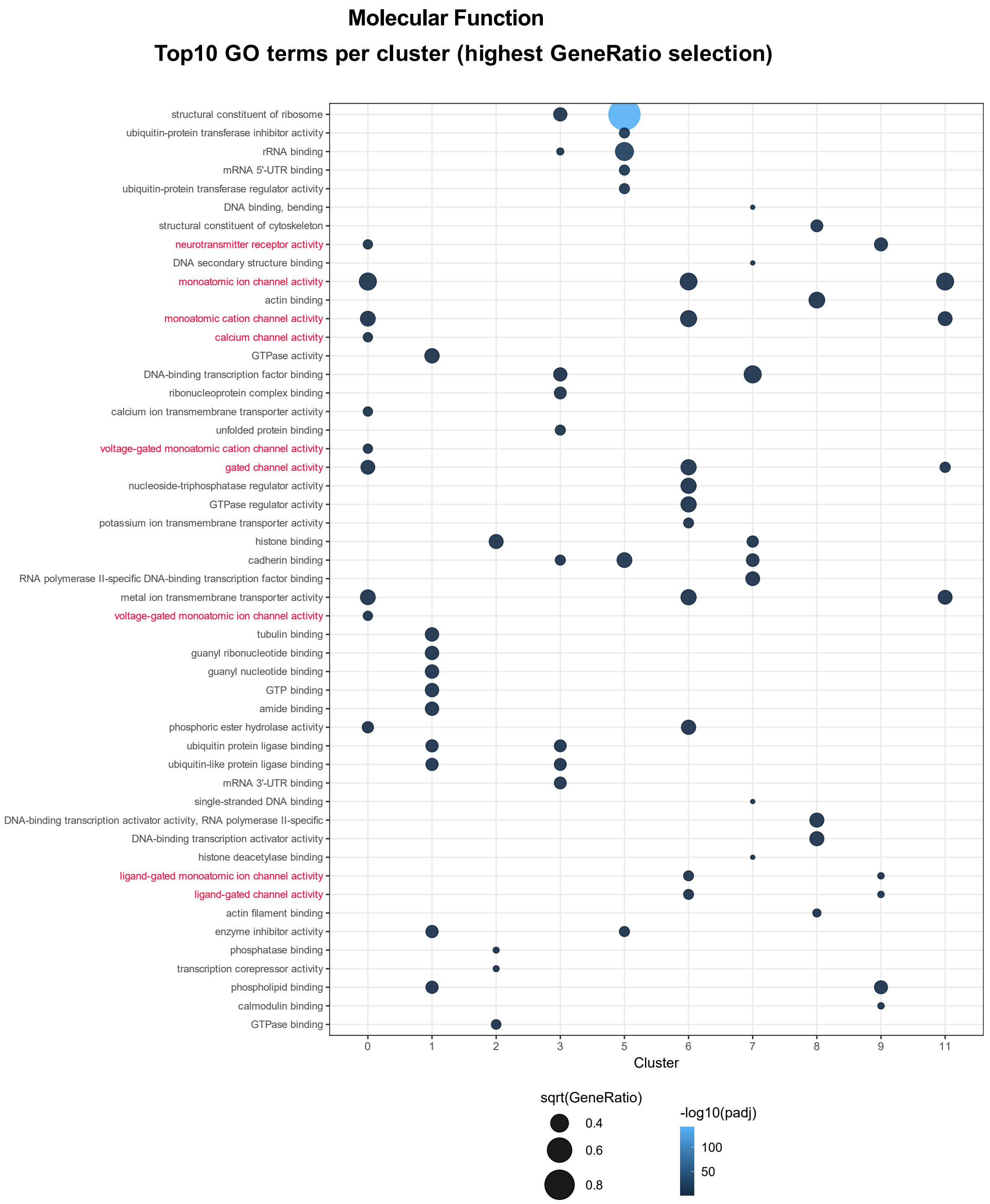

#### Supplemental Information 1

A. **DNA sequences used for production of GST-Npvf and GST-Vgll2 fusion proteins in *E. coli*. for antibody generation. Sequences cloned into vector pGEX-4T3.**

**Npvf**

gaattcATGGAAATCATCTCCCTGAAACGCTTTATTCTGCTGACGGTTGCCACGAGTAGCTTCC
TGACCTCTAACACCTTCTGTACGGACGAATTTATGATGCCGCATTTCCACAGCAAAGAAGG
CGATGGTAAATATTCTCAGCTGCGTGGCATTCCGAAAGGTGAAAAAGAACGCAGTGTTTCC
TTTCAAGAACTGAAAGACTGGGGCGCAAAAAACGTGATCAAAATGAGTCCGGCGCCGGCC
AACAAAGTTCCGCATTCCGCGGCCAATCTGCCGCTGCGTTTCGGTCGCACCATTGATGAAA
AACGTAGTCCGGCAGCTCGCGTCAATATGGAAGCGGGCACGCGTTCACACTTTCCGTCGCT
GCCGCAGCGTTTCGGTCGTACCACGGCACGCTCACCGAAAACCCCGGCTGACCTGCCGCAA
AAACCGCTGCATTCGCTGGGCAGCTCTGAACTGCTGTACGTCATGATTTGCCAGCACCAAG
AAATCCAGTCTCCGGGCGGTAAACGTACCCGTCGCGGTGCATTTGTGGAAACGGATGACGC
CGAACGCAAACCGGAAAAAtaatgatagctcgag

**Vgll2**

gaattcATGAGCTGCCTGGATGTTATGTACCAAGTGTATGGCCCGCCGCAACCGTATTTCGCGG
CGGCGTATACCCCGTATCACCAAAAACTGGCGTACTATAGCAAGATGCAGGAAGCGCAGG
AATGCGCGAGCCCGGGTAGCAGCGCGAGCGGTAGCAGCAGCTTCAGCAACCCGACCCCGG
CGAGCGTGAAAGAGGAAGAGGGTAGCCCGGAAAAAGAGCGTCCGCCGGAAGCGGAGTAC
ATCAACAGCCGTTGCGTTCTGTTCACCTATTTTCAGGGCGACATTAGCAGCGTGGTTGATGA
ACACTTCAGCCGTGCGCTGAGCCACCCGAGCAGCTACACCCCGAGCTGCACCAGCAGCAA
AGCGCATCGTAGCAGCGGTCCGTGGCGTGCGGAGGGCACCTTTCCGATGAGCCAGCGTAGC
TTCCCGGCGAGCTTTTGGAACAGCGCGTACCAAGCGCCGGTGCCGGCGCCGCTGGGTAGCC
CGCTGGCTGCGGCGCACAGCGAACTGCCGTTTGCGACCGACCCGTATAGCCCGGCGACCCT
GCACGGTCACCTGCACCAGGGTGCGGCGGATTGGCACCACGCGCACCCGCACCATGCGCA
CCCGCACCATCCGTACGCGCTGGGTGGCGCGCTGGGCGCGCAAGCGAGCGCGTATCCGCGT
CCGGCGGTGCATGAGGTTTATGCGCCGCACTTCGACCCGCGTTATGGTCCGCTGCTGATGC
CGGCTGCGACCGGTCGTCCGGGCCGTCTGGCGCCGGCGAGCGCGCCGGCGCCGGGCAGCC
CGCCGTGCGAACTGGCGGCGAAGGGCGAGCCGGCGGGCAGCGCGTGGGCGGCGCCGGGTG
GCCCGTTTGTGAGCCCGACCGGTGACGTTGCGCAAAGCCTGGGTCTGAGCGTTGATAGCGG
CAAACGTCGTCGTGAATGCAGCCTGCCGAGCGCGCCGCCGGCGCTGTATCCGACCCTGGGC
taatgatagctcgag

B. **DNA sequences used to misexpress human VGLL2 and TEAD1 in human hypothalamic organoids**

**VGLL2**

ATGAGCTGTCTGGATGTTATGTACCAAGTCTATGGTCCTCCGCAGCCCTACTTCGCA
GCCGCCTACACCCCCTACCACCAGAAACTAGCCTATTATTCCAAAATGCAGGAAGC
GCAGGAGTGCAATGCCAGCCCCAGCAGCAGTGGCAGCGGCAGCTCCTCATTTTCCA
GCCAAACCCCAGCCAGTATAAAAGAGGAAGAAGGCAGCCCAGAGAAAGAGCGCCC
ACCAGAGGCAGAGTACATCAACTCCCGCTGCGTCCTCTTCACTTATTTCCAGGGGGA
CATCAGCTCCGTGGTGGATGAACATTTCAGCAGGGCCCTGAGCCAACCCAGCAGCT
ACTCTCCTAGCTGTACCAGCAGCAAAGCACCAAGGAGCTCTGGGCCCTGGCGAGAC
TGCTCCTTCCCGATGAGCCAGCGCAGCTTCCCCGCCTCCTTCTGGAATAGCGCGTAC
CAGGCGCCAGTGCCCCCGCCGCTGGGCAGCCCTCTGGCCACCGCGCACTCGGAGCT
GCCCTTCGCCGCCGCCGACCCCTACTCGCCCGCCGCGCTGCATGGCCACCTGCACCA
GGGCGCCACGGAGCCCTGGCACCACGCGCACCCGCACCACGCGCACCCGCATCACC
CCTACGCCCTGGGCGGCGCCCTCGGCGCCCAGGCCGCCCCCTACCCGCGCCCCGCCG
CCGTGCACGAAGTCTACGCGCCGCACTTCGACCCGCGCTATGGGCCGCTGCTGATGC
CAGCCGCCTCGGGGCGCCCGGCCCGCCTCGCAACCGCCCCGGCGCCCGCGCCCGGC
AGTCCTCCCTGCGAGCTCTCCGGCAAAGGCGAGCCGGCGGGCGCCGCGTGGGCCGG
GCCCGGGGGACCCTTCGCGAGCCCCTCGGGGGACGTGGCCCAGGGTCTGGGCCTCA
GCGTGGACTCAGCTCGTCGTTATTCCCTCTGTGGTGCATCCCTCCTGAGCTGA

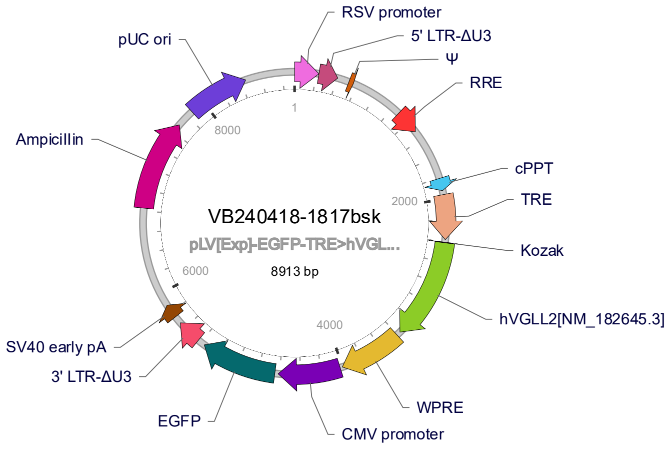


**TEAD1**

ATGGAGCCCAGCAGCTGGAGCGGCAGTGAGAGCCCTGCCGAAAACATGGAAAGGA
TGAGTGACTCTGCAGATAAGCCAATTGACAATGATGCAGAAGGGGTCTGGAGCCCC
GACATCGAGCAAAGCTTTCAGGAGGCCCTGGCTATCTATCCACCATGTGGGAGGAG
GAAAATCATCTTATCAGACGAAGGCAAAATGTATGGTAGGAATGAATTGATAGCCA
GATACATCAAACTCAGGACAGGCAAGACGAGGACCAGAAAACAGGTGTCTAGTCAC
ATTCAGGTTCTTGCCAGAAGGAAATCTCGTGATTTTCATTCCAAGCTAAAGGATCAG
ACTGCAAAGGATAAGGCCCTGCAGCACATGGCGGCCATGTCCTCAGCCCAGATCGT
CTCGGCCACTGCCATTCATAACAAGCTGGGGCTGCCTGGGATTCCACGCCCGACCTT
CCCAGGGGCGCCGGGGTTCTGGCCGGGAATGATTCAAACAGGGCAGCCAGGATCCT
CACAAGACGTCAAGCCTTTTGTGCAGCAGGCCTACCCCATCCAGCCAGCGGTCACA
GCCCCCATTCCAGGGTTTGAGCCTGCATCGGCCCCAGCTCCCTCAGTCCCTGCCTGG

CAAGGTCGCTCCATTGGCACAACCAAGCTTCGCCTGGTGGAATTTTCAGCTTTTCTC
GAGCAGCAGCGAGACCCAGACTCGTACAACAAACACCTCTTCGTGCACATTGGGCA
TGCCAACCATTCTTACAGTGACCCATTGCTTGAATCAGTGGACATTCGTCAGATTTAT
GACAAATTTCCTGAAAAGAAAGGTGGCTTAAAGGAACTGTTTGGAAAGGGCCCTCA
AAATGCCTTCTTCCTCGTAAAATTCTGGGCTGATTTAAACTGCAATATTCAAGATGA
TGCTGGGGCTTTTTATGGTGTAACCAGTCAGTACGAGAGTTCTGAAAATATGACAGT
CACCTGTTCCACCAAAGTTTGCTCCTTTGGGAAGCAAGTAGTAGAAAAAGTAGAGA
CGGAGTATGCAAGGTTTGAGAATGGCCGATTTGTATACCGAATAAACCGCTCCCCAA
TGTGTGAATATATGATCAACTTCATCCACAAGCTCAAACACTTACCAGAGAAATATA
TGATGAACAGTGTTTTGGAAAACTTCACAATTTTATTGGTGGTAACAAACAGGGATA
CACAAGAAACTCTACTCTGCATGGCCTGTGTGTTTGAAGTTTCAAATAGTGAACACG
GAGCACAACATCATATTTACAGGCTTGTAAAGGACTGA

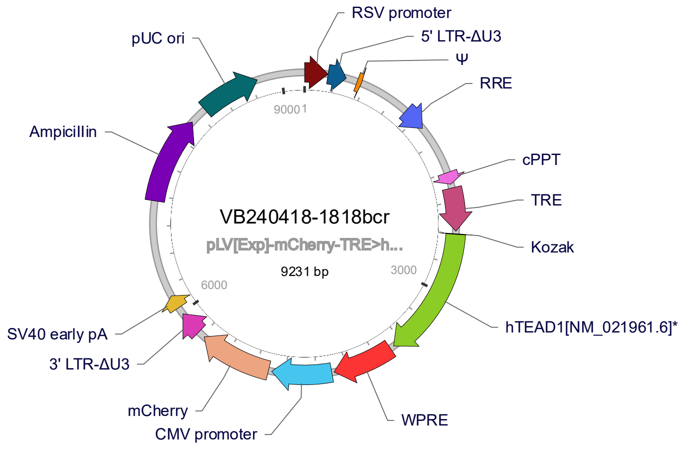
